# The structural landscape of Microprocessor mediated pri-*let-7* miRNA processing

**DOI:** 10.1101/2024.05.09.593372

**Authors:** Ankur Garg, Renfu Shang, Todor Cvetanovic, Eric C. Lai, Leemor Joshua-Tor

**Affiliations:** W. M. Keck Structural Biology Laboratory, Cold Spring Harbor Laboratory, One Bungtown Road, Cold Spring Harbor, New York, 11724 USA; Howard Hughes Medical Institute, Cold Spring Harbor laboratory, One Bungtown Road, Cold Spring Harbor, New York, 11724 USA; Developmental Biology Program, Sloan Kettering Institute, 430 East 67th St, ROC-10, New York, NY 10065, USA

## Abstract

miRNA biogenesis is initiated upon cleavage of a primary miRNA (pri-miRNA) hairpin by the Microprocessor (MP), composed of the Drosha RNase III enzyme and its partner DGCR8. Multiple pri-miRNA sequence motifs affect MP recognition, fidelity, and efficiency. Here, we performed cryo-EM and biochemical studies of several let-7 family pri-miRNAs in complex with human MP. We show that MP has the structural plasticity to accommodate a range of pri-miRNAs. These structures revealed key features of the 5’ UG sequence motif, more comprehensively represented as the “fUN” motif. Our analysis explains how cleavage of class-II pri-let-7 members harboring a bulged nucleotide generates a noncanonical precursor with a 1-nt 3’ overhang. Finally, the MP-SRSF3-pri-let-7f1 structure reveals how SRSF3 contributes to MP fidelity by interacting with the CNNC-motif and Drosha’s PAZ-like domain. Overall, this study sheds light on the mechanisms for flexible recognition, accurate cleavage, and regulated processing of different pri-miRNAs by MP.

## Introduction

MicroRNAs (miRNAs) are ∼ 22 nucleotides (nt) non-coding RNAs that generally repress mRNA expression post-transcriptionally^1^. miRNAs are processed from hairpin-containing transcripts called primary miRNAs (pri-miRNAs), typically transcribed by RNA polymerase II. In animals, the canonical pri-miRNA stem-loop is cleaved by the Microprocessor (MP), a heterotrimeric nuclear complex containing the RNase III endonuclease Drosha, and two copies of its essential co-factor, DGCR8. MP cleavage results in a staggered cut in the pri-miRNA to produce a shorter hairpin precursor (pre)-miRNA of ∼ 70-80 nt,^2–4^ which is exported to the cytoplasm by exportin 5^5^. The cytoplasmic RNase III endonuclease, Dicer, cleaves off the apical loop of the pre-miRNA to generate a ∼ 22 base-pair (bp) miRNA duplex^6^. This duplex is loaded into an Argonaute (Ago) protein, where one strand, the guide RNA, remains in the complex forming the mature RNA-induced Silencing Complex (RISC)^7^. RISC is then guided to the target RNA, via base-pairing, predominantly through the miRNA seed sequence (nt ∼2-8)^8,9^, which ultimately results in target repression.

Apart from its role in initiating miRNA processing, MP acts as a gatekeeper that permits only specific hairpin transcripts, out of an ocean of plausible miRNA-like structures, to enter the biogenesis pathway. The most prominent features are single-stranded regions flanking a ∼ 35 ± 1 bp double-stranded stem often harboring several wobbles, mismatched base pairs, bulges, and a ≥ 10 nt apical loop^10–14^. In addition, pri-miRNAs are enriched for certain short motifs. These motifs affect cleavage efficiency and fidelity of MP-mediated pri-miRNA processing (M^2^P^2^) and include a basal UG dinucleotide motif at the -14 nt position in the 5’-strand, a UGUG motif at the 3’ end of the 5’ arm (at the base of the apical loop), a CNNC (where N is any nucleotide) motif at the -17 nt position in the 3’-strand and a GHG (where H is not G) mismatch (mGHG) in the 3’-strand at the -3 nt to -5 nt position in the lower stem region^13,15–18^ (Figure 1A). All these motifs are recognized directly by the MP^13,19^, with the exception of the 3’ CNNC-motif, which recruits SRp20/SRSF3 to assist MP in the productive processing of pri-miRNAs^15,20^. Nonetheless, pri-miRNAs vary considerably in their structures. This is especially pronounced in the wide variety of sequences, shapes, and sizes of the apical loop, even within a given family of pri-miRNAs, such as the pri-let-7 family^21^. In addition, a large fraction of miRNAs appears to lack all these motifs. Recent cryo-EM studies provided a glimpse into how the 5’ UG^22^ and mGHG^23^ motifs are recognized by the MP. However, the features dictating MP recognition of the large pool of pri-miRNAs lacking these motifs, and the plasticity the MP must have to accommodate the large variation in pri-miRNA structures, remain poorly understood.

**Figure 1.**
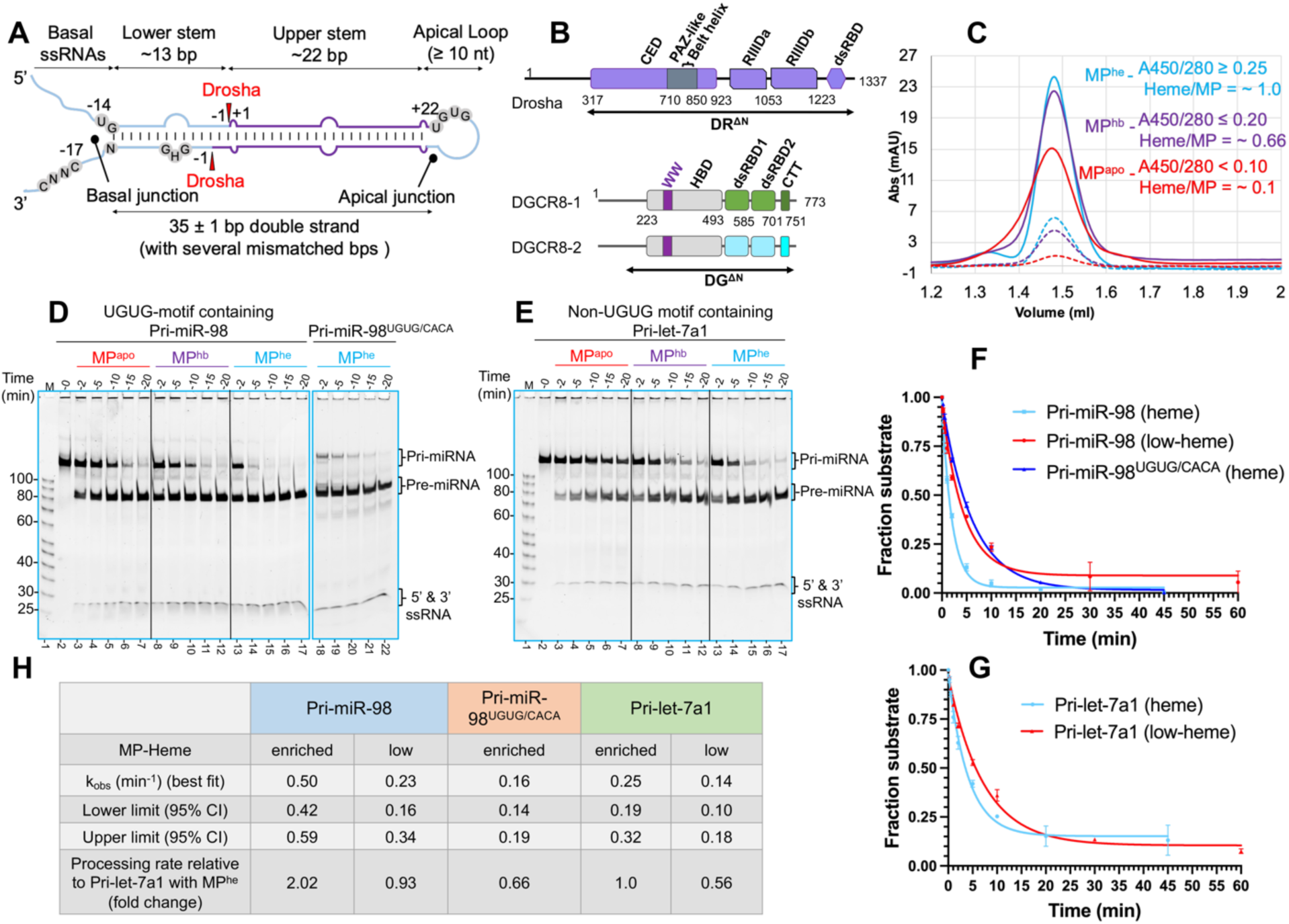
Heme has a ubiquitous role in M^2^P^2^. (A) General architecture of a canonical pri-miRNA. The short sequence motifs and Drosha cleavage sites are shown. (B) Domain architecture of human Drosha (DR) and DGCR8 (DG). The DR^ΔN^ and DG^ΔN^ truncations used in this study are underlined. (C) Analytical SEC analysis for different MP heme-variants: MP^he^, MP^hb^ & MP^apo^. The UV280 and UV450 traces are shown in solid and dotted lines, respectively. Average A450/280 ratios and calculated heme/MP molar ratios are shown. (D, E) Qualitative cleavage assays, (F, G) plots for RNA substrate disappearance in M^2^P^2^ in near-pre-steady state, and (H) the calculated cleavage rates for pri-miR-98 and (E) pri-let-7a1 by MP^apo^, MP^hb^ and MP^he^.

The 3’ CNNC-motif is the most conserved sequence motif, present in ∼ 60% of human pri-miRNAs, and is specifically bound by a critical splicing factor, SRSF3/SRp20^15,20,24^. In addition to rescuing the productive M^2^P^2^ of 201 human pri-miRNAs (∼11% of the total pri-miRNA pool) in an *in-vitro* cleavage assay, SRSF3 modulates alternative processing events and suppresses unproductive/abortive processing (nick processing and inverse processing) for hundreds of pri-miRNAs^18^. Although SRSF3 assists Drosha recruitment at the basal junction for an accurate RNA cleavage^20^, a mechanistic view of SRSF3’s effect on pri-miRNA processing remains unknown.

Approximately 26% of human pri-miRNAs contain the apical UGUG motif^15^, and their M^2^P^2^ fidelity is dependent on heme binding at the WW motif (heme-dimerization motif) in the heme-binding domain (HBD) of DGCR8 (Figure 1B)^25–28^. Heme facilitates HBD dimerization^29^, and selectively impacts the M^2^P^2^ of pri-miRNAs^30^, but heme’s role in MP biology is still unclear, especially for non-UGUG type pri-miRNAs.

One of the first miRNA families discovered, the let-7 miRNAs, contains 12 functionally conserved members in humans^31^. MP processing categorizes these pre-let-7s into two distinct classes: three pre-let-7s belong to class-I with the canonical 2-nt 3’overhang, and nine belong to class-II with just 1-nt overhang at the 3’ end^21^. Class-II pre-let-7s are mono-uridylated by a Terminal Uridylyl transferase-(TUT)4/7^32–34^ converting them to a suitable Dicer substrate, with a 2-nt 3’ overhang, ensuring their efficient processing into the mature miRNAs^35^. A bulged nucleotide at the 5’ cleavage site in class-II pri-let-7s is thought to be responsible for their non-canonical M^2^P^232^. However, the molecular basis for this difference in processing is still unclear.

In this study, we took a structural and biochemical approach to understand how the MP handles the wide range of pri-miRNA substrates, expand our understanding of MP motif recognition, and uncover how SRSF3 modulates the M^2^P^2^ for CNNC-containing pri-miRNAs.

## Results

### Heme facilitates M^2^P^2^

To study the effect of heme binding to the MP complex on pri-miRNA processing, we expressed and purified an N-terminally truncated MP (DR^ΔN^DG^ΔN^) (Figure 1B) in low-heme (MP^apo^) and endogenous heme-bound (MP^hb^) forms (Figure S1A). By supplementing the cell culture media with 5-aminolevulinic acid (5-ALA), we expressed and purified a heme-enriched MP (MP^he^) (Figure S1B). The average UV 450/280 absorbance (A450/280) ratio for purified MP^he^, MP^hb^ and MP^apo^ was ≥ 0.25, ≤ 0.20 and < 0.10, respectively (Figure 1C). The calculated “heme/MP” molar ratio for MP^he^, MP^hb^ and MP^apo^ were ∼ 1, ∼ 0.66 and ∼ 0.10, respectively, indicating differential heme occupancies for these purified MP complexes.

We subjected these to an *in-vitro* time-course cleavage assay using pri-miR-98, a UGUG-motif containing pri-let-7, as a substrate. We observed that MP^he^ processing of pri-miR-98 into pre-miR-98 is significantly faster than, either with MP^hb^ or MP^apo^, indicating that MP activity on the UGUG-motif containing pri-miR-98 substrate is heme-dependent (Figure 1D). Mutation of the UGUG motif in pri-miR-98 to CACA, results in slower processing by MP^he^, suggesting that the UGUG-motif plays a role in M^2^P^2^ of pri-miR-98 (Figure 1D compare lane 13-17 to 18-22). Interestingly, similar RNA processing differences were observed for pri-let-7a1 (a non-UGUG motif containing pri-miRNA) (Figure 1E). A quantitative analysis for pri-miR-98 and pri-let-7a1 processing also shows a clear reduction in substrate cleavage rate with MP^apo^ compared to MP^he^ (Figure 1F, 1G), while it is reduced for the pri-miR-98^UGUG/CACA^ mutant even with MP^he^ (Figure 1F, H). Cleavage of different pri-let-7’s by MP^he^ and MP^apo^ also exhibited significant variability in qualitative (Figure S1C-D) and quantitative cleavage assays (Figure S1E-H), demonstrating heme’s role in all pri-miRNA processing, likely linked to reduced flexibility in MP upon heme-binding.

### The pri-let-7s-MP^he^ complex reveals new features (Class-II let-7s)

To understand how the MP accommodates the large variation in pri-let-7 pri-miRNAs we determined cryogenic electron microscopy (cryo-EM) structures of human MP^he^ in complex with three pri-let-7s in their pre-catalytic state (Figure 2A). For these studies, we used Drosha isoform 4, DR^ΔN^ (aa 317-1337), and DGCR8 DG^ΔN^ (aa 175-751) (Figure 1B). We will be using Drosha isoform 1 numbering for consistency. The overall resolutions for the cryo-EM maps were 3.2 Å, 3.3 Å and 2.9 Å for pri-miR-98, pri-let-7a1 and pri-let-7f1 bound to MP^he^, respectively, as estimated from their GSFSC curves (Figure S2A, S2D, S2G). Although local resolution varies for these structures, Drosha and most of the pri-miRNAs were observed at high resolution for all three, while DGCR8 and the apical loop of the pri-miRNA were at lower resolution (Figure S2B, S2E, S2H).

**Figure 2.**
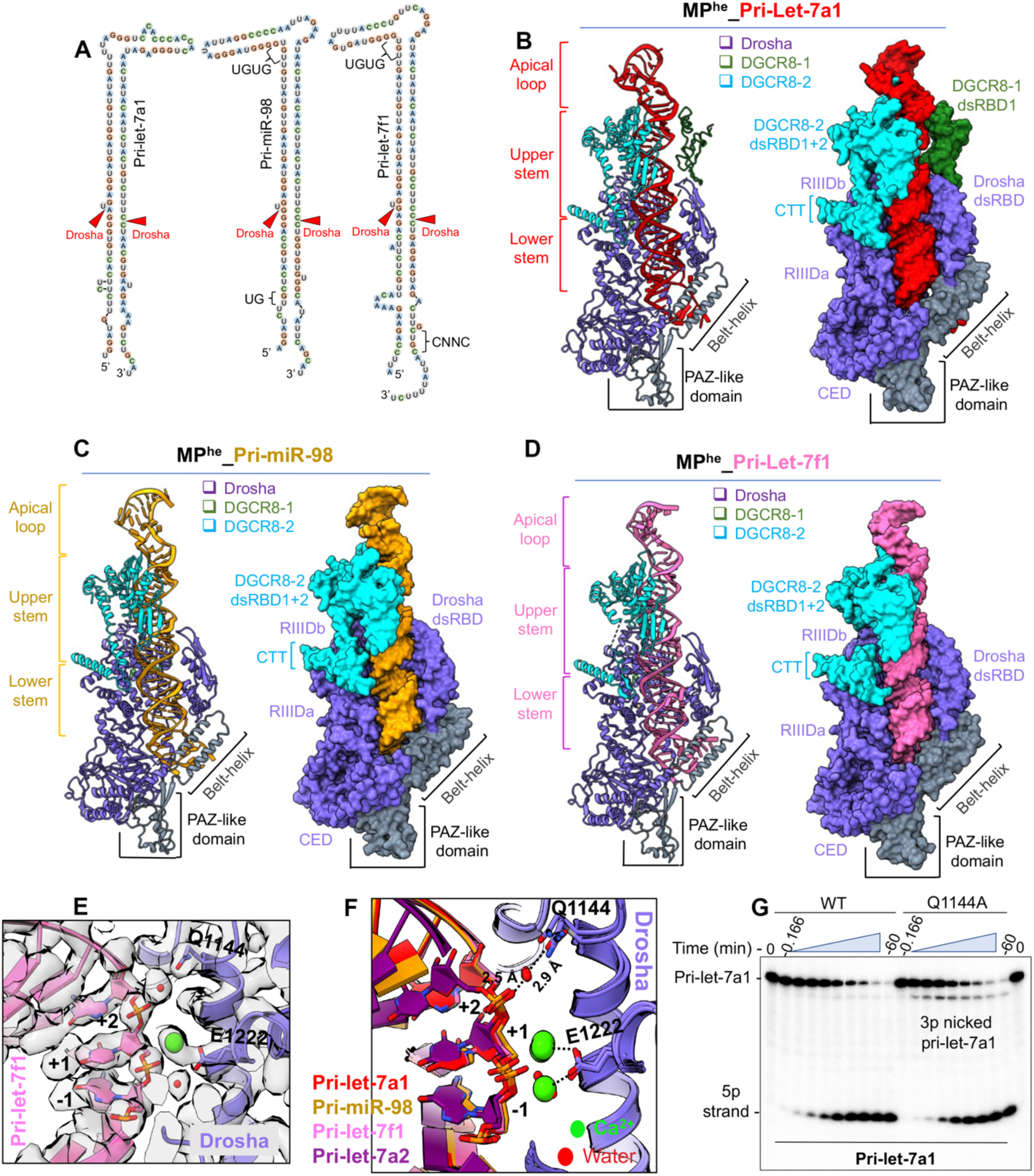
Cryo-EM structure of MP^he^ in complex with class-II pri-let-7s. (A) 2D schematics of different class-II pri-let-7s used for cryoEM. Conserved sequence motifs and Drosha cleavage sites are shown. (B) Cartoon and space fill representation of the cryo-EM structures of MP^he^-pri-let-7a1, (C) MP^he^-pri-miR-98, and (D) MP^he^-pri-let-7f1 complexes. (E) A zoom-in view of the 5’ catalytic site in MP^he^-pri-let-7f1 map showing cryoEM density of the Ca^2+^ ion (green) and water molecules (red), (F) the conserved interactions in multiple pri-let-7 structures stabilizing the 5p +2 geometry via Q1144. (G) Pri-let-7a1 processing assay with MP^he^-Q1144A mutant showing accumulation of 3’ nicked RNA products compared to MP^he^.

Overall, our MP^he^-pri-let-7 structures display the familiar canonical heterotrimeric arrangement of the Drosha-(DGCR8)_2_ complex (Figure 2B-D)^19,22,23^ trapped in a pre-catalytic state. The heterotrimer is stabilized by the interaction of Drosha RIIIDa and RIIIDb with the CTT peptides from DGCR8-2 and DGCR8-1, respectively. Both dsRBDs from DGCR8-2 in all three structures (Figure 2B-D), and DGCR8-1’s dsRBD1 could be confidently built in the MP^he^-pri-let-7a1 (Figure 2B), but not in other structures to avoid overfitting into a low-quality portion of the map. Cryo-EM density was not observed for DGCR8-1 dsRBD2, which is likely flexible due to the absence of stabilizing interactions. All cryo-EM maps show a significant chunk of density above the DGCR8-dsRBDs, which corresponds to the RNA apical loop and the DGCR8 HBD dimer (Figure S3A-B). Cryo-EM map denoising allowed us to visualize partial secondary structure features in this density, but due to the lack of any guiding structure for this region, we could not build the complete HBD into the map. In addition, AlphaFold^36,37^ prediction models do not match the observed density (Figure S3C-D). Nonetheless, the crystal structure of the dimerization subdomain, comprised of the WW motif (PDBid 3LE4)^28^ could be confidently docked into the map away from the RNA. The WW motif presents the interaction surface for heme and includes two cysteines that act as ligands to the heme iron^25,29^, though the heme itself was not resolved in our structures nor in the crystal structure. Interestingly, this subdomain has no direct contact with the RNA (Figure S3A-B).

In all three structures, the pri-let-7 stem adopts a near A-form conformation over ∼ 3.2 helical turns, covering ∼ 35 base pairs. As observed in previous studies, the dsRNA-ssRNA junction (called the basal junction) is clamped between an α-helical hairpin (the ‘belt’) emanating from the PAZ-like domain, and the wedge loop (amino acids 930-952) from Drosha^22,23^. Of this, two turns of the RNA double-helix (∼ 22 bp) are docked onto the central groove formed between the two RNase III domains (RIIID) and dsRBD of Drosha, while ∼ 1.2 turns (∼ 13 bp) are engulfed by the DGCR8 dsRBDs (Figure 2B-D). Thus, the RNA in the pre-catalytic state is stabilized via interactions with all three proteins of the complex all along the RNA backbone.

The 5’ and 3’ RNA cleavage sites for the different pri-let-7s are positioned at the catalytic sites of RIIIDb and RIIIDa, respectively. A Ca^2+^ ion, used in place of the Mg^2+^ ion to prevent catalysis, is accommodated in either one or both RIIIDs in these structures. However, the observed density for the Ca^2+^ ion in RIIIDb is consistently stronger than at the RIIIDa site, likely due to the differential binding of Ca^2+^ ions in the two RIIIDs, as was observed previously^23^. Additionally, our cryo-EM maps revealed a conserved water molecule stabilizing the 5p +2 nt phosphate via Gln1144 from RIIIDb, anchoring the RNA backbone and rigidifying the downstream 5’ cleavage site (Figure 2E). This water-mediated interaction is observed in all our structures (Figure 2F and Figure S3E-H). Mutation of Gln1144 to an alanine affects 5p strand cleavage for pri-let-7a1, generating significantly more 3’ nicked RNA products (Figure 2G). In the 5’ cleavage site, Glu1222 is known to coordinate the catalytic Mg^2+^ ion^16,19^ and our study shows that Gln1144-mediated 5p +2 phosphate stabilization is also important for efficient pri-miRNA cleavage.

Our cryo-EM structures now show the complete 35 to 36 bp RNA stem (Figure 2B-D), a hallmark of canonical pri-miRNAs^12,13,38^. These structures confirm that the pri-let-7 stem can accommodate many non-Watson-Crick (WC) bps at different positions, without major perturbations in the RNA helix. The cryo-EM maps exhibited helical density atop the RNA stem region (Figure S3A-B) and aided by the crystal structure of the Lin28-pre-let-7 complex (PDBid 5UDZ)^39^, we were able to model this density as the part of the apical loop in different pri-let-7s. However, the nucleotides corresponding to the unpaired loop region could be built for pri-let-7a1 structure only (Figure 2B). The UGUG motif in pri-miR-98 and pri-let-7f1 (Figure S3J-K) and the Lin28-interacting GGAG/GAAG motif in pri-let-7a1 (Figure S3L)^40,41^, both in the apical loop, could also be confidently traced. The apical loop in each of the three pri-let-7s adopts a hairpin structure with ∼ ½ turn of ds-helical geometry, dominated by G-C base-pairing, which along with the unpaired loop nucleotides, is exposed in all three MP^he^-bound structures, and might serve as a point of contact for other RNA-binding proteins (RBPs) to modulate M^2^P^2^ activity. Interestingly, the UGUG motif is solvent exposed, while the GGAG/GAAG motif abuts the unmodelled HBD density.

### RNA drives MP domain repositioning for pri-let-7 cleavage

To highlight the structural differences in class-I and class-II pri-let-7 binding in MP, we determined a 2.9 Å Cryo-EM structure of MP^he^ with class-I pri-let-7a2 (Figure S4A-B) in the pre-catalytic state, and compared it with our class-II pri-let-7 structures described above.

The MP^he^-pri-let-7a2 cryo-EM structure shows Drosha, positioned on the RNA basal junction, with its dsRBD interacting with the RNA upper stem region (Figure 3A). Notably, both DGCR8-CTTs interact with Drosha, though in this structure cryo-EM density corresponding to the DGCR8-HBD-dsRBDs and a portion of pri-let-7a2’s upper stem and apical loop was not well resolved (Figure S4C). Drosha is positioned almost identically on the RNA basal junction, with an almost identical lower stem RNA geometry in all the pri-let-7s structures described here (Figure 3B), as well as in the structure of the complex with pri-miR-16-2^23^. However, the RNA upper stem in class-I pri-let-7a2 is bent compared to the class-II pri-let-7s (Figure S4D). Similarly, class-I pri-miR-16-2^23^ exhibits a similar bend of the upper stem relative to the Drosha-lower stem RNA complex (Figure 3C). Notably, the kink in the helix starts right after the 5’ cleavage site, which results in a lateral displacement of up to 10 Å and 21 Å of the 3p and 5p nucleotides at the +20^th^ base-pair position in the upper stem, respectively (Figure 3C).

**Figure 3.**
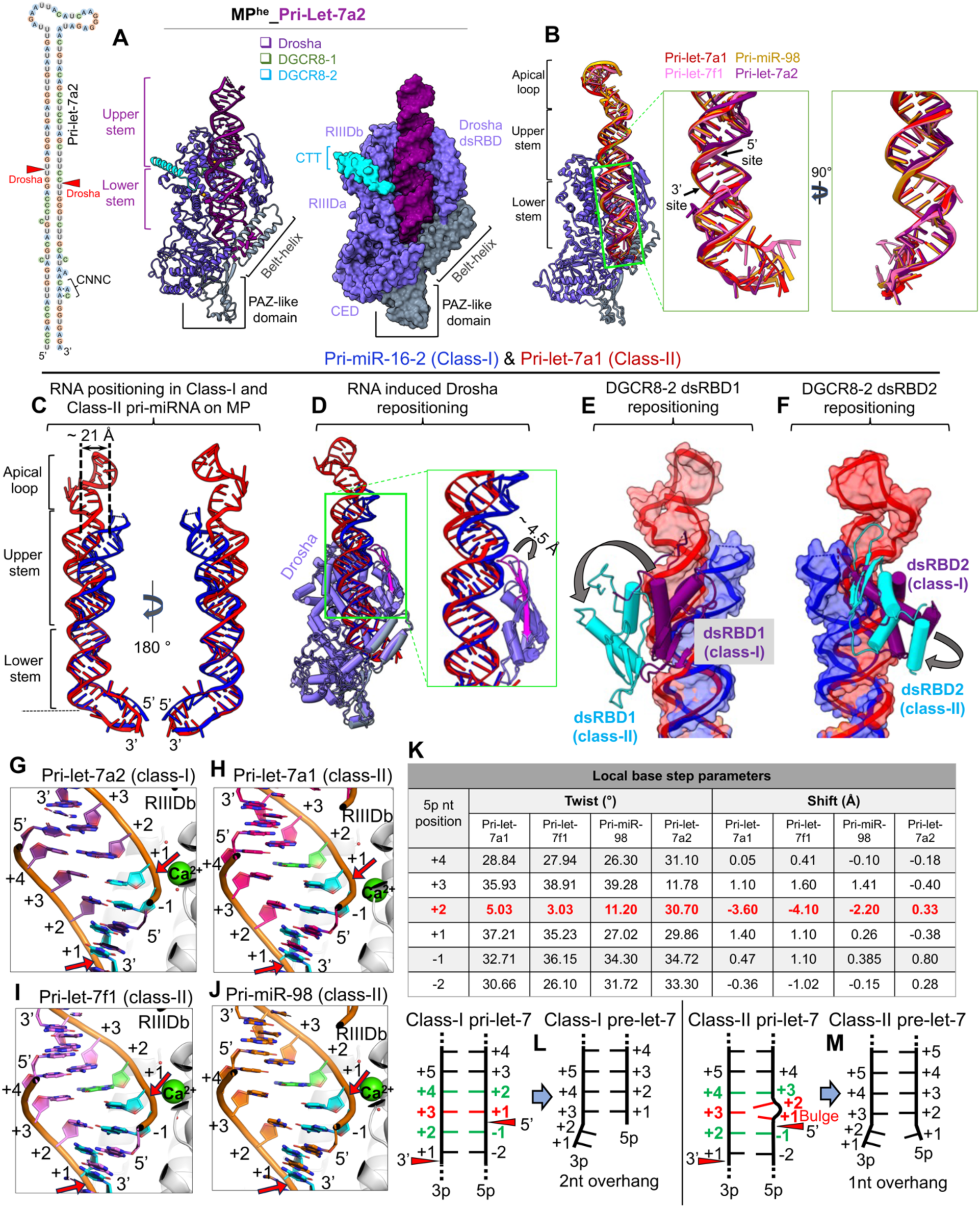
RNA-driven structural rearrangements in MP in the two pri-let-7 classes. (A) 2D schematics of the pri-let-7a2, and the cryo-EM structure of the MP^he^-pri-let-7a2 complex. (B) Drosha superimposition showing the conserved positioning of the RNA lower stem in different pri-let-7s. The difference in the (C) RNA positioning (D) Drosha dsRBD, (E) DGCR8-2 dsRBD1 and (F) DGCR8-2 dsRBD2 positioning in class-I (pri-miR-16-2 in blue) and class-II (pri-let-7a1 in red-orange) pri-miRNA bound MP^he^ structures. The repositioning in different components are shown with arrows and distances are also marked. Base-pairing patterns at the 5’ cleavage site in (G) class-I pri-let-7a2, and class-II (H) pri-let-7a1 (I) pri-let-7f1 and (J) pri-miR-98. The 5p +1nt (cyan) in pri-let-7a2 is paired with 3p +3nt, while it remains unpaired/bulged in class-II structures. Cleavage sites are highlighted with red arrows. (K) The local base twist and shift parameters for the 5p nucleotides around the 5’ cleavage site. The 5p +2 shows distorted geometry (highlighted in red) accommodating for the bulged 5p +1 nt. A schematic of the base-pairing patterns observed in class-I pre-let-7 (L) and class-II pri-let-7s (M) when bound to MP in their pre-catalytic state. Drosha cleavage sites are shown with red arrows. The bulged 5p +1 nt in class-II pri-let-7s stays unpaired and changes the upstream base-pairing register, allowing MP to generate pre-let-7 species with a 1nt 3’ overhang.

Bending of the RNA upper stem is complemented by structural rearrangements in both Drosha and DGCR8. The Drosha dsRBD’s β-hairpin (β1-β2) opens ∼4.5 Å from the of the domain to accommodate the movement of the RNA upper stem in class-I pri-miRNA (pri-miR-16-2 in the figure) (Figure-3D). All three observed dsRBDs spatially reposition themselves with the RNA upper stem in class-II pri-let-7a1 compared to class-I pri-miR-16-2 (Figure-3E-F) such that all the RNA-specific interactions are conserved in both class-I and class-II pri-miRs. This indicates that the MP undergoes an RNA-induced conformational rearrangement, accommodating the changes in RNA structure. Taken together, the different cryo-EM structures reveal the plasticity of MP’s dsRBDs to recognize and incorporate different pri-let-7 miRNAs, driven by rearrangements in the RNA upper stem.

### Generation of class-I vs. class-II pre-miRNAs

To understand how the bulged nucleotide at 5p +1 position in class-II pri-let-7 processing assists MP in generating the two different 3’ overhangs, we compared the two classes of pri-let-7 RNAs around the cleavage sites. In class-I pri-let-7a2 the 3p +2, +3 and +4 nts pair up with the 5p -1, +1, and +2 nts, respectively (Figure 3G). For class-II structures, the pairing between bases are shifted by one nucleotide after the bulge (Figure 3H-J). The 5p +2 nt and its pairing with the 3p +3 nt appear to absorb most of the distortion from the presence of the bulge immediately preceding it (Fig 3K, S4F-G), with its backbone bulging outward from the RNA helix (Figure S4E). This geometry allows the RNA duplex to accommodate the bulge. The bulge stacks on top of the 5p -1 base and is on a plane with the 3p +3 base without base pairing with it (Figure 3H,3J). The consequence of these acrobatics is that it “straightens out” the overall trajectory of the RNA upper stem (see above and Figure 3C) and positions the unpaired 5p +1nt above the catalytic ion adding an extra nucleotide in 5p strand, thus leaving a single nucleotide as the 3’ overhang in class-II pre-let-7s (Figure 3L-M).

### The UG or fUN motif

The UG motif at the 5p -14nt position is prevalent in ∼25% of human pri-miRNAs, and is tightly enclosed by Drosha’s belt, the wedge loop, and its dsRBD. The high resolution of the cryoEM maps in this region in our structures allowed us to confidently analyze features of UG motif. The U in the MP^he^-pri-miR-98 structure is flipped out with its base stacking onto His802 from one of the helices of the Drosha belt. The base identity is recognized through H-bonds between O2(U) and O4(U) and Arg1273 (from Drosha dsRBD) and Ile942 main chain amide (from Drosha wedge), respectively (Figure 4A). Its phosphate interacts with Arg938 (from the wedge loop). The following G in pri-miR-98, at position -13, pairs with a G on the 3’ strand via its Hoogsteen edge establishing the first base pair in the lower stem (Figure 4A). The only other UG motif containing pri-miRNA structure, pri-miR-16-1 (PDBid 6LXD) also exhibits similar interactions for this motif except that -13 G pairs with a C on the 3p strand through canonical Watson-Crick hydrogen bonding^22^ (Figure S4H).

**Figure 4.**
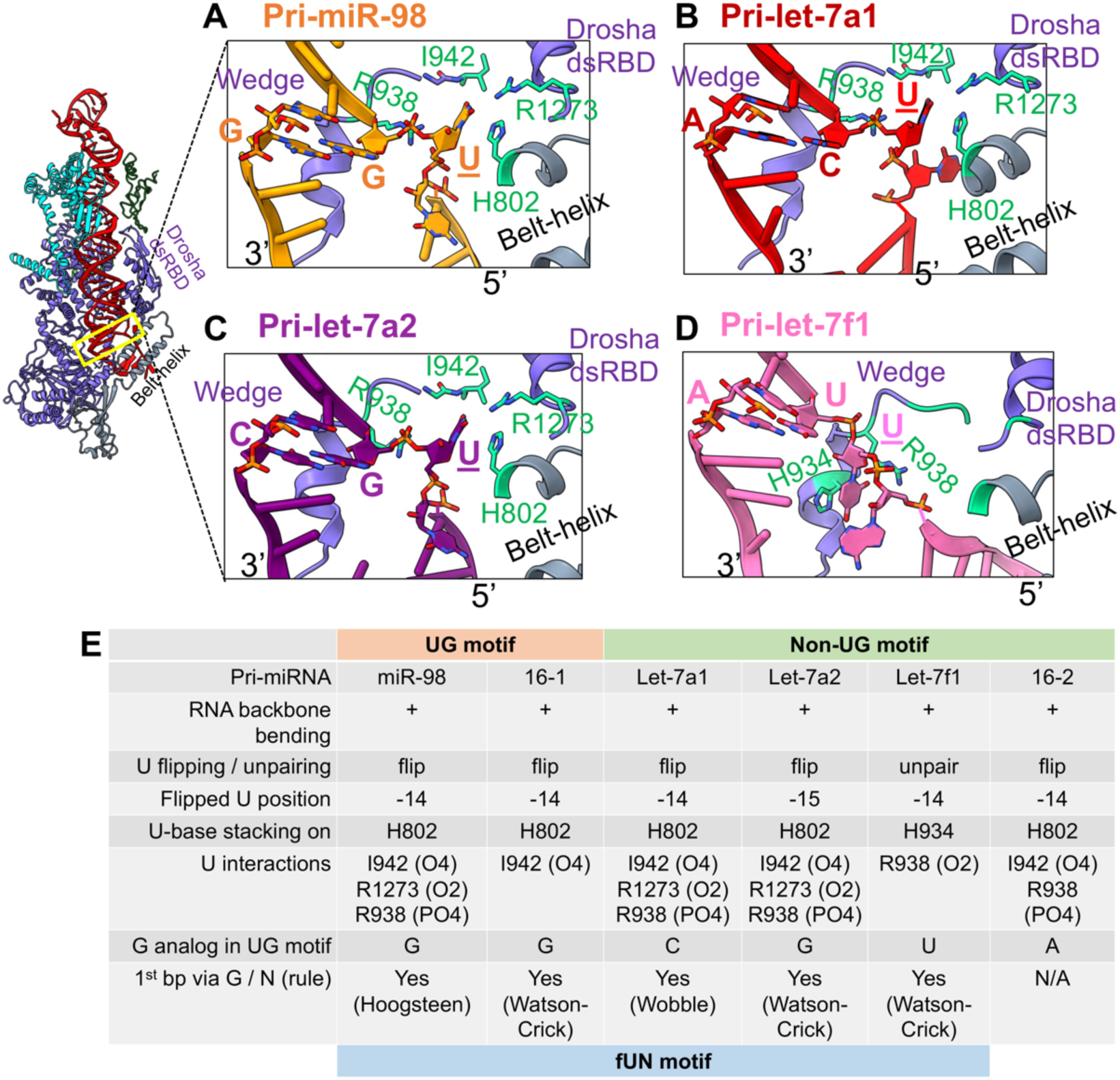
The 5’ UG or fUN motif? A zoomed-in view of the interactions observed in the UG (or fUN) motif in (A) pri-miR-98, (B) pri-let-7a1, (C) pri-let-7a2 and (D) pri-let-7f1 with residues (green sticks) from, Drosha’s wedge, dsRBD, and the belt-helix. The U flips out (underscored) while N establishes the 1^st^ base-pairs in the RNA lower stem. (E) Conserved structural features characteristic of this motif.

Many pri-miRNAs lack the UG motif *per se*; however, more than 90 other pri-miRNAs have a U at the 5p -14nt position. Three of our structures, the MP^he^ complexes of pri-let-7a1, pri-let-7a2, and pri-let-7f1 do not contain a UG motif, but all have an unpaired U at the -14nt position, or -15nt position (pri-let-7a2), and in all but one (pri-let-7f1), this U is flipped out (Figure 4A-D). For the pri-let-7f1 complex, the U base stacks onto His934 (from the Drosha wedge); it is the O2 atom that interacts with Arg938 (from the wedge loop) rather than its phosphate, as in the other structures. Similar to the pri-miRNAs with a UG motif, the following nucleotide in all these structures pairs with a nucleotide from the 3’ arm to form the 1^st^ base-pair of the lower stem, either through canonical Watson-Crick (Figure 4C-D) or via non-canonical base pairing (Figure 4A-B). In the structure with pri-miR-16-2 RNA (PDBid 6V5B) the -14 U is flipped out as well, but the following nucleotide stays unpaired (Figure S4I).

Overall, our structural analysis shows that whether the pri-miRNA has a strict UG motif or not, the U at position -14 is flipped or is at least unpaired and is sensed by several amino acids from Drosha, while the -13 position nucleotide forms the first base pair in the RNA lower stem, whether it is a G or not (Figure 4E), though a strong G enrichment at the -13 position was reported previously^15^. We, therefore, propose to revise the ‘UG sequence motif’ designation to a ‘flipped U with paired N’ (fUN) structural motif positioned predominantly at the 5p -14 position. The fUN motif would allow a more comprehensive classification of the pri-miRNA pool.

### SRSF3-assisted M^2^P^2^ via the 3’ CNNC motif

To demonstrate the role of SRSF3 in M^2^P^2^, we purified and included recombinant human SRSF3 in RNA processing assays. *In-vitro* processing of CNNC motif-containing pri-let-7c (CGUC) by MP^he^ primarily results in unproductive cleavage and minimal pre-let-7c product (productive cleavage) (Figure 5A lane 1-6 and Figure S1C lane 8-13). Thus, any improvement in pre-let-7c yield could be reliably observed and quantified. Adding SRSF3-FL to the RNA cleavage reaction significantly improved productive cleavage of pri-let-7c (Figure 5A: compare lanes 8-12 to lanes 2-6 and Figure S5B), while with the RRM domain alone (aa 1-84) much less pre-let-7c is produced (Figure S5C lane9-13). Furthermore, a CNNC mutation to UUUU resulted in loss of SRSF3-mediated enhancement of pri-let-7c^CNNC/UUUU^ productive cleavage (Figure 5A lane14-19). In near-pre-steady state conditions, both the rate of pri-let-7c substrate cleavage and the rate of inverse processing exhibit a subtle reduction as SRSF3 concentration is increased from a sub to supra-stoichiometric ratio relative to the RNA (Figure 5B, 5C, 5G). However, the extent of the inverse-cleavage products decreases (Figure 5C), while the pre-let-7c product increases (Figure 5D) with increasing SRSF3 concentrations. These observations clearly highlight SRSF3’s role in M^2^P^2^.

**Figure 5.**
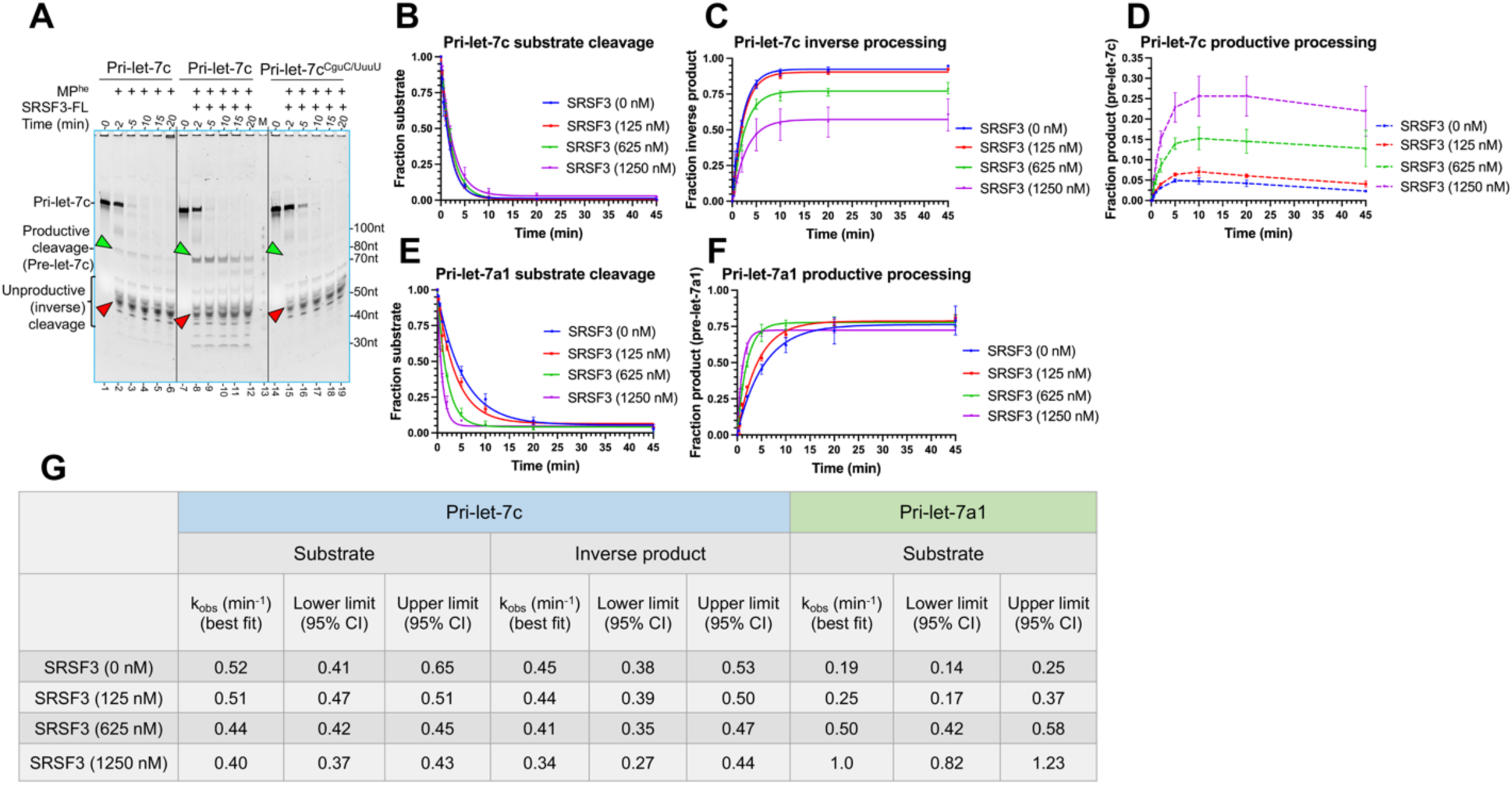
SRSF3 assists Drosha processing of pri-let-7s. (A) Qualitative cleavage assays of pri-let-7c and pri-let-7c^CNNC/UUUU^ by MP^he^, showing the effect of SRSF3 on pre-let-7c product formation (green arrowhead) and unproductive cleavage products (red arrowhead). (B) Plot for pri-let-7c substrate disappearance (C) inverse processing and (D) pre-let-7c appearance in near-pre-steady state, showing the effect of SRSF3 in M^2^P^2^. As formed pre-let-7c product decreases overtime, its traces are shown as connected lines only. (E) Plot for pri-let-7a1 cleavage and (F) pre-let-7a1 appearance in near-pre-steady state, showing SRSF3 effect on M^2^P^2^. (G) The calculated rates for pri-let-7s processing with SRSF3.

Additionally, for the CNNC-motif containing pri-let-7g, the effects on M^2^P^2^ are better observed in the unproductive cleavage products, which are reduced when incubated with SRSF3 (Figure S5D: compare lane 2-6 with lane 8-12). The substrate cleavage rate is drastically improved up to 8-fold in near-pre-steady state conditions (Figure S5F-G). The impact of SRSF3 on substrate cleavage rate and unproductive cleavage products is completely abolished with mutation of the CNNC-motif (Figure S5D lane 14-19, S5F-G) within pri-let-7g, further underscoring the importance of the CNNC-motif. Interestingly, for the CNNC-deficient pri-let-7a1, substrate catalysis seems positively impacted in qualitative cleavage assays (Figure S5E). While the cleavage rate increases up to 5-fold with increasing SRSF3 concentrations (Figure 5E, 5F, 5G), the extent of product production doesn’t change much, suggesting that SRSF3 might have a general role in M^2^P^2^.

### The cryoEM structure of the MP^he^-pri-let-7f1-SRSF3 complex

To understand how SRSF3 ensures M^2^P^2^ fidelity, we determined a 3.1 Å cryo-EM structure of the quaternary complex containing MP^he^-pri-let-7f1 with SRSF3-FL assembled in a pre-catalytic state (Figure S6A-E). The cryo-EM map revealed unambiguous density not only for the MP and the majority of the RNA, but also for SRSF3 and the extended 3p RNA strand containing CNNC-motif (Figure S6B-D) near Drosha’s PAZ-like domain and the belt.

The MP^he^-pri-let-7f1-SRSF3 structure shows Drosha and DGCR8 interacting with the basal junction and upper stem, respectively. Both dsRBDs and the CTT of DGCR8-1, and only the CTT for DGCR8-2 was observed (Figure 6A-B) in the structure, while the HBD could not be modeled in the low-resolution cryoEM density for this region (Figure S6C-E). In this structure, the 3p strand is resolved to position 22, which includes the complete CNNC-motif, while only 17nts were well-ordered in the MP^he^-pri-let-7f1 ternary structure. Though SRSF3-FL was used for the cryo-EM sample preparation, we could clearly position and rebuild the NMR structure of RRM domain (PDBid 2I2Y)^42^ and build the linker peptide (thumb peptide) of the RS domain (aa 1-87) (Figure 6B), but density for the remainder of the RS domain (aa 88-164) was not observed. Nevertheless, the local resolution of SRSF3 and the RNA in that region is 3.0 – 4.0 Å (Figure S6E). The MP^he^-pri-let-7f1 geometry is largely unchanged upon SRSF3 binding, which in turn grabs onto the CNNC motif in the extended 3p RNA strand, like a hand gripping a rope, and nestles onto the preformed Drosha PAZ-like domain surface, burying ∼703 Å^2^ surface area to form a new protein-protein interface (Figure 6A-B). The Drosha PAZ-like domain exhibits only minor structural rearrangements upon SRSF3 binding with an RMSD of 0.26 Å over 70 Cα atoms.

**Figure 6.**
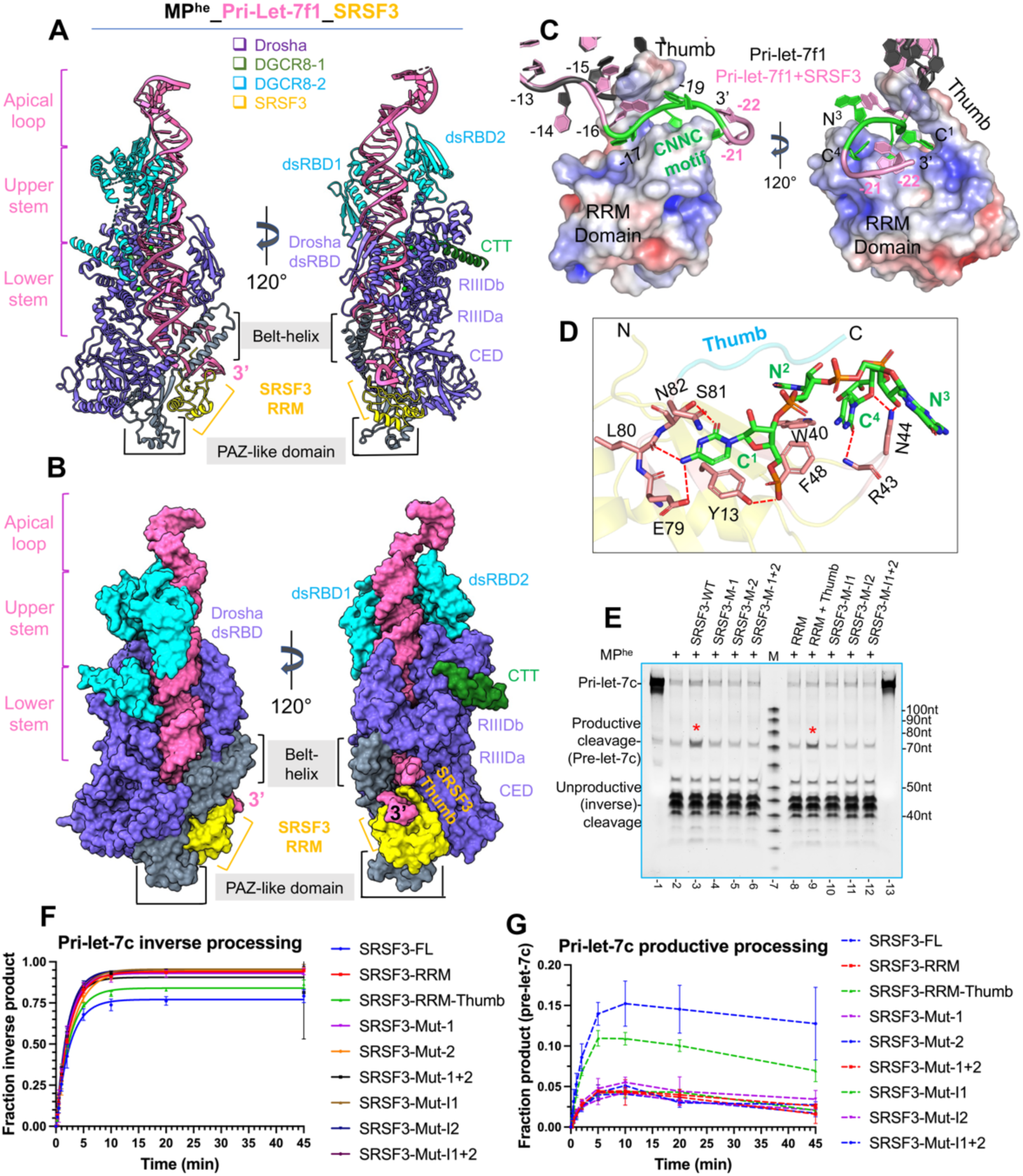
The MP^he^-pri-let-7f1-SRSF3 quaternary complex. (A) Cryo-EM structure of MP^he^-pri-let-7f1-SRSF3 in the pre-catalytic state shown in cartoon and (B) surface representation. The SRSF3 RRM domain (yellow) is bound to 3p CNNC motif in pri-let-7f1 (pink) and is docked onto Drosha’s PAZ-like domain (grey). The thumb peptide is clearly visible and interleaved between the two RNA strands. (C) The pri-let-7f1 3p ssRNA region in the MP^he^ (dark grey) and MP^he^-SRSF3 structures (pink) shown in two orientations. The 3p strand in the MP^he^ structure (dark grey) would clash with the SRSF3 thumb peptide. In the MP^he^-SRSF3 bound structure, the CNNC motif (green) passes through the electrostatically charged channel formed by the SRSF3 RRM domain and thumb. Mapping of the electrostatics on the SRSF3 surface is shown (-5 (red) to +5 kT (blue)). (D) Molecular interactions between the CNNC motif (green sticks) and SRSF3 RRM domain (beige-colored sticks). Direct H-bonding interactions are shown as red dotted lines. (E) *In-vitro* M^2^P^2^ of pri-let-7c with different SRSF3 mutations/truncations. Only the RRM-thumb restores pre-let-7c (marked) levels to SRSF3-WT level. (F) Plot for the pri-let-7c inverse processing and (G) pre-let-7c formation in near-pre-steady state, showing the effect of SRSF3 mutations/truncations in M^2^P^2^. As formed pre-let-7c product decreases overtime, its traces are shown as connected lines only. SRSF3-M-1, M-2, and M-1+2 indicate SRSF3 with mutations in C^1^, C^4^, or C^1^+C^4^ nucleotide-stabilizing residues in the CNNC motif, respectively. RRM and RRM thumb denotes SRSF3^1-84^ and SRSF3^1-90^ truncations. SRSF3-M-I1, M-I2, and M-I1+2 are SRSF3 with mutations in the Drosha interface 1, 2, or 1+2, respectively.

### CNNC motif recognition

The SRSF3 RRM domain adopts a canonical βαββαβ fold^43^ (Figure S7A) exposing several hydrophobic residues in its β-sheet (Figure S7C), while the linker peptide (aa 84-87) folds on top of it to create a positively charged channel for RNA binding. SRSF3 clamps the CNNC-motif (also denoted as 3p -17 C^1^N^2^N^3^C^4^ motif below) between its “thumb” (the linker peptide) and ‘fingers’ (the RRM β-sheet) (Figure 6C and Figure S7B).

The SRSF3 thumb is inserted between the two RNA strands, gripping the 3p strand, ensuring that the 3p nucleotides -15 through -18 do not base pair with the 5p strand, as has been observed for pri-let-7f1, pri-miR-98 and elsewhere^22^. Arg86 from the thumb is inserted between the -15 and -16 bases, and distorts the RNA backbone, positioning the -17 C^1^ base in the RNA binding channel on SRSF3 (Figure 6C). The C^1^ base is stacked between Asn82 and Tyr13 (from β1) and establishes multiple H-bonds with Glu79, Ser81 side chains and the Leu80 backbone oxygen. Also, the C^1^ phosphate H-bonds with Tyr13 (Figure 6D). The N^2^ base (U in pri-let-7f1) undergoes stacking interactions with Trp40 (from β2), and N^3^ (G in pri-let-7f1) flips outwards and binds to Asn44 via its sugar 2’-OH. Additionally, the flipped nucleotide base points into a negatively charged shallow groove on the Drosha belt-helix (formed by Glu822, Glu823, and Gln826). The RNA backbone at C^4^ kinks, which flips its base underneath the N^2^-N^3^ backbone and docks it into a positively charged shallow groove in the SRSF3 RRM domain formed by Trp40, Asn44, and Phe48, and forms a H-bond between O2(C^4^) and the main chain amine of Arg43. Moreover, its sugar O1’ forms a H-bond with Asn44 (Figure 6D and S7C), stabilizing the kinked geometry. Beyond the CNNC motif, the RNA backbone further bends and stacks the -21 A base onto the N^2^ (U) base (Figure S7A-B). The -22 U base stacks onto the -21 A base without any direct contact with SRSF3 or Drosha.

Comparison of the SRSF3-CNNC component in the MP^he^ bound structure (Figure S7A-C) with the NMR structure of the SRSF3^RRM^-rCAUC tetranucleotide complex^42^ (PDBid 2I2Y) (Figure S7D-F) revealed distinctive features in pri-miRNA recognition by both the RRM and thumb peptide, highlighting a context-specific RNA recognition by SRSF3.

Mutation of SRSF3 residues interacting with either C^1^, C^4^, or both (Figure S5A), lead to much less pre-let-7c in qualitative assays (Figure 6E lane 3-6) and in near-pre-steady state conditions (Figure 6F, 6G). Moreover, the SRSF3 RRM domain alone produces much less pre-let-7c, but adding the thumb increases pre-let-7c level close to FL-SRSF3 levels both, in qualitative (Figure 6E lane 3, 8-9) and in near-pre-steady state conditions (Figure 6F, 6G). These observations validate the role of both C^1^ and C^4^ nucleotide recognition by SRSF3 RRM and the thumb for an effective CNNC motif-mediated response. However, the substrate cleavage rate for the pri-let-7c shows no significant changes for any of these mutations/truncations compared to WT-SRSF3 (Figure S5H, S5I).

### The SRSF3 – Drosha interface

The MP-SRSF3-pri-let-7f1 structure revealed a new protein-protein interface between SRSF3 and Drosha’s PAZ-like domain. The PAZ-like domain is essentially unchanged upon SRSF3 binding, implying that its surface is pre-formed for SRSF3 interaction. We should note, however, that this surface did change upon pri-miRNA binding. At the SRSF3-PAZ-like domain interface, the PAZ-like domain exhibits a negative electrostatic potential on one side of its surface and a deep positively charged groove on the other, at the 3p RNA exit site (Figure S7G). Notably, the SRSF3 RRM exhibits the opposite charged surfaces at its interface with the Drosha PAZ-like domain (Figure S7H) and docks perfectly onto that surface (Figure 7A). Therefore, there is both charge and surface complementarity between the two proteins.

**Figure 7.**
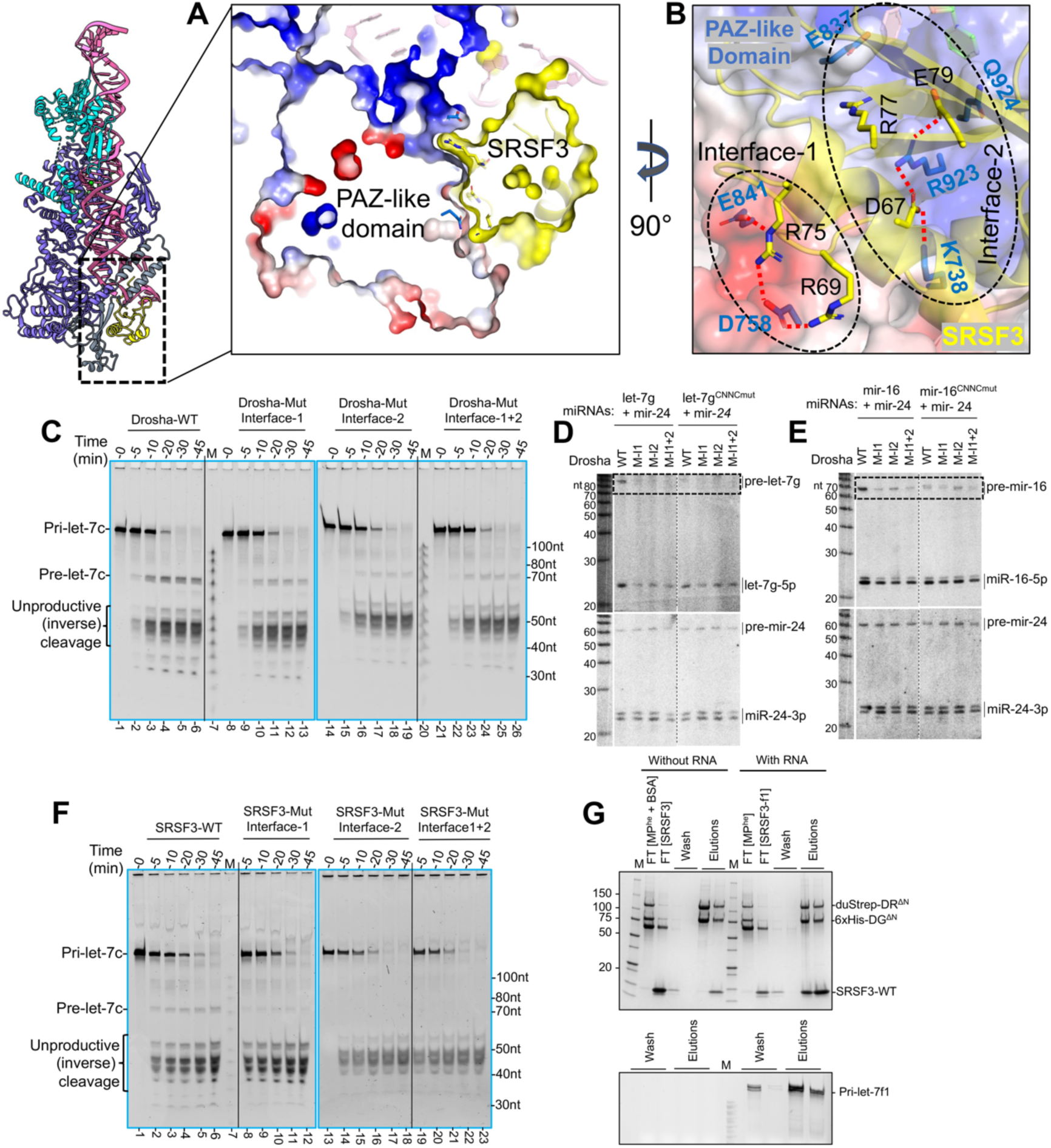
The SRSF3-Drosha PAZ-like domain interface. (A) A cutaway view of the Drosha-SRSF3 interface showing SRSF3 (yellow) perfectly nestled into the PAZ-like domain scaffold (shown as an electrostatic surface). (B) The interaction between SRSF3 (yellow sticks) and the PAZ-like domain (sky blue sticks), clusters into two regions, interface-1 and interface-2 (black dotted ovals) with complementary charges between the two proteins. (C) *In-vitro* processing of pri-let-7c using Drosha-mutations in the Drosha-SRSF3 interface. Mutations in Interface-1, 2 or 1+2 show less or no pre-let-7c product during the time course. (D-E) Northern analysis of miRNA processing in HEK293T *Drosha-KO* cells rescued by wt or mutant Drosha constructs. Cells were co-transfected with expression constructs for CNNC-bearing pri-let-7g (D) or pri-miR-16 (E), or counterparts with mutations in CNNC motifs; non-CNNC pri-miR-24 serves as a negative control. Mutations in the SRSF3-interacting interface of Drosha reduce pri-miRNA processing for CNNC-bearing miRNAs, as clearly indicated by the reduction of pre-miRNA hairpin species (dotted boxes). These Drosha mutants also exhibit impaired accumulation of mature let-7g and miR-16 (asterisks), with a stronger effect on let-7g. Biogenesis of non-CNNC pri-miRNA constructs was not affected by Drosha mutations. (F) *In-vitro* M^2^P^2^ of pri-let-7c with SRSF3-mutants at the Drosha-SRSF3 interface. (G) SRSF3 pull-down assay with MP^he^ and pri-let-7f1. SRSF3-WT co-elutes with the MP^he^-pri-let-7f1 as analyzed on SDS-PAGE (upper panel) and urea-PAGE (lower panel) gels.

The positive protrusion on SRSF3 establishes a series of salt bridges with residues from the PAZ-like domain, which we call “Interface-1**”**. For “Interface-2”, negative residues from SRSF3 form another set of salt bridges with residues from the PAZ-like domain and wedge-loop. Additionally, SRSF3 Glu79 is stacked against Gln924 from the Drosha wedge. SRSF3 Arg77 is stacked with Glu837 from the PAZ-like domain and points into a charged pocket near the 3’ RNA exit site (Figure 7B).

To validate these observations, we mutated interface-1 and interface-2 residues in the two proteins independently, to oppositely charged residues and analyzed them in pri-let-7c cleavage assays. Purified SRSF3 R69E/R75E (interface-1), D67H/E79R/R77A (interface-2) mutants (Figure S5A), and Drosha D758H/E841R (interface-1), K738E/R923E/Q924A (interface-2) mutants eluted at the same retention volume as their respective WT proteins, indicating no protein folding issues. Compared to WT MP^he^, the interface1 or interface-2 Drosha MP^he^ mutants result in significantly less pre-let-7c product formation in qualitative cleavage assay (Figure 7C).

We further analyzed mutations in the SRSF3-interacting interface of Drosha and the requirement for pri-miRNA CNNC in cells. We rescued MP activity in *Drosha-KO* HEK293T cells by introducing wt or mutant Drosha constructs. At the same time, we co-transfected wt or CNNC-mutated miRNA constructs (let-7g or mir-16), along with the non-CNNC mir-24 as a control. Northern blotting showed that all three Drosha mutations compromised pri-miRNA processing, specifically for CNNC-containing miRNAs. This was most clearly seen by the reduced levels of pre-let-7g and pre-miR-16; mature let-7g was also strongly reduced while accumulation of mature miR-16 was impaired (Figure 7D-E). It is possible that the perdurance of mature miRNAs obscures the effect of reduced nuclear cleavage of pri-miR-16. By contrast, CNNC-mutant pri-let-7g and pri-miR-16, along with non-CNNC pri-mir-24, were all equivalently processed by wt and Drosha variants bearing mutations at the SRSF3 interface (Figure 7D-E).

The *in-vitro* pri-miRNA binding for these mutated MP^he^ is largely unaffected compared to WT-MP^he^, (Figure S7I-J), suggesting that the observed effect is not due to impaired RNA binding. Similarly, compared to the WT-SRSF3, the SRSF3 mutations in interface-1 or interface-2 produced almost no pre-let-7c product in qualitative cleavage assays (Figure 7F), and failed to induce a quantitative effect on inverse-cleavage product (Figure 6F) or pre-let-7c (Figure 6G) in near-pre-steady-state conditions. However, the pri-let-7c cleavage rate is not impacted compared to WT-SRSF3 (Figure S5H, S5I). To discern whether the reduced M^2^P^2^ of pri-let-7c in SRSF3 mutants is a consequence of their impaired Drosha PAZ-like domain docking or an impaired RNA interaction, we performed *in-vitro* pull-down assays using the MP^he^-RNA as bait and gel-shift assay using pri-let-7 miRNAs. We found that the WT-SRSF3 co-elutes with the MP^he^-RNA complex (Figure 7G) and binds with the pri-miRNAs (Figure-S7K). Notably, both SRSF3 interface-1 and interface-2 mutants fail to co-elute with the MP^he^-RNA complex (Figure-S7L-N), while only the interface-2 mutant still binds the pri-miRNAs (Figure-S7K). These results suggest that the reduced M^2^P^2^ of pri-let-7c in interface-2 mutation is a direct consequence of impaired PAZ-domain docking. Overall, the mutation analysis supports the structural observation, that the interface between PAZ and SRSF3 is crucial for productive processing of pri-miRNAs.

## Discussion

Given the breadth of miRNA-mediated silencing in regulating gene expression across eukaryotes, accurate and efficient miRNA biogenesis is of premier interest. We have long been intrigued by the structural diversity of pri-miRNA substrates that the MP can tackle, even within the same miRNA family. Using the most abundant miRNA family in the human genome, the let-7 family, we investigated how MP recognizes and processes these diverse pri-miRNAs into pre-miRNAs.

In preparing the MP complex, we found that proper incorporation of heme significantly increases the substrate cleavage rate whether or not a UGUG motif, previously implicated in heme recognition, is present in the apical loop of pri-miRNA. Our structures showed that the heme-binding region, the WW motif of DGCR8, was quite distant from the RNA and the UGUG motif on the apical loop, but might contribute to MP activity by promoting dimerization of DGCR8-HBD, which could affect its interface with the pri-miRNA apical loop. In addition, Heme didn’t appear to have an effect on proper loading of the MP on the substrate in our studies, in contrast to previous suggestions^30^.

We were able to observe the almost complete pri-miRNA, revealing the relative position of the apical-loop (preE) on top of the dsRNA stem (Figure 2B-D), in addition to the UGUG-motif, the GGAG-motif and the well-studied CNNC-motif (Figure 6). The pri-let-7’s apical loop in the different structures is exposed at least on one surface and could serve as a binding site for other RNA binding proteins (RBPs) to influence the M^2^P^2^ of specific pri-miRNAs, including Syncrip^44^, Musashi-1^45^, or Lin28B^46^. Superimposing the crystal structure of Lin28a-let-7f1 preE (PDBid 5UDZ)^39^ onto the MP^he^-pri-let-7f1 structure reported here revealed that the zinc-knuckle of Lin28 would clash with DGCR8 dsRBD’s binding to pri-let-7 (Figure-S3I) and might negatively impact M^2^P^2^ of pri-let-7s. While, the Lin28 cold-shock domain (CSD) would be positioned on the ss-loop away from DGCR8.

Let-7 pri-miRNAs segregate into two distinct classes based on whether a canonical 2-nt overhang is produced at the 3’-end for class-I pre-let-7s, or a single nt is left as the 3’ overhang in the class-II variety. It has been long speculated that a bulge at the 5p cleavage site is responsible for this difference. Here we show that the bulge alters the overall trajectory of the upper stem, essentially ‘straightening out’ the pri-miRNA (Figure 3C, S4D) and perturbs the local RNA strand geometry (Figure 3H-J, S4E-G). This alters the pairing register in the upper stem, resulting in a 1nt 3’ overhang following the cleavage reaction (Figure 3L-M). Quite often, a protein induces structural changes on nucleic acids upon binding^47,48^, but in M^2^P^2^ the RNA drives the repositioning of several domains of the enzyme as observed in class-I and class-II pri-let-7 structures. (Figure 3D-F).

Importantly, we addressed two characteristic sequence motifs in our study. The first is the UG motif at the 5p -14 position, which we suggest renaming the fUN (flipped U with paired N) motif. We find that the U is unpaired in all cases we examined and is usually flipped (Figure 4, S4H-I). The role of the following nucleotide is to establish the first base pair in the lower stem. Though G is more prevalent at this position in metazoan pri-miRNAs, and was shown to independently contribute to RNA catalysis^15^, it appears to vary. The positioning of this motif is slightly flexible and can be similarly accommodated at position -15.

Second, the CNNC-motif at the 3p -17 position has a role in productive M^2^P^2^, mediated by the splicing factor SRSF3, which modulates at least 84% of Drosha-dependent miRNAs^18^. Our study shows how SRSF3 grabs onto the CNNC motif like a hand grabbing a rope (Figure 6A-B), between the thumb and the RRM domain (Figure 6C). It is also tucked onto Drosha with complementary surfaces and electrostatics (Figure 7A-B). Thus, the MP-pri-miRNA complex is in a conformation that can bind SRSF3 without changes. SRSF3 situates Drosha firmly on the basal junction and prevents the pairing of the ss regions of the two strands, which might extend the ds lower stem, as seen in pri-miR-16-1^22^, causing aberrant loading of the MP on the pri-miRNA.

SRSF proteins usually bind their target RNA through their RRM domain, while the RS-domain participates in protein-protein interaction^49,50^. Though some RRM domains are known to directly interact with proteins via their α1 helix and β1-3 sheet^51–53^, SRSF3 RRM uses its α2-β4 side for Drosha interaction. The SRSF3 thumb is known to interact with NXF1/TAP during mRNA export^42^, and the same region interacts with the CNNC in M^2^P^2^, exhibiting its multifunctional nature. Another splicing factor, the SR protein SRSF7 is also known to be involved in Drosha recruitment to the basal junction^15,24^. Its RRM shares ∼73% identity with the SRSF3, with all residues involved in CNNC binding and Drosha PAZ-like domain interactions conserved, suggesting a similar molecular mechanism for SRSF7.

Interestingly, MP^he^ pulls down SRSF3 in the absence of RNA (Figure 7G) and has a different sensitivity *in vitro* to the mutants we examined in the presence of RNA (Figure S7L-N). Since the Drosha surface that interacts with SRSF3 changes upon pri-miRNA binding^23^, this implies that SRSF3 binds to a different surface of Drosha in the absence of RNA.

The MP-pri-mRNA structures reported here and elsewhere reveal MP’s invariable interactions with the lower stem of the RNA, measuring the distance from the basal junction to the cleavage sites. The presence of conserved sequence motifs in the lower stem does not appear to affect Drosha-lower stem geometry. This rigidity is complemented by its plasticity to reorient its dsRBDs with the upper stems of pri-miRNAs to accommodate a wide landscape of pri-miRNA structures. The combination of a rigid lower-stem length and interface, with the ability to reorient modules that could bind various characteristic motifs helps the MP distinguish these substrates from other structured RNAs.

## Acknowledgments

We thank members of the Joshua-Tor laboratory for helpful discussions. We thank CSHL Cryo-EM facility and the CSHL mass Spectrometry shared resources. We thank V. N. Kim (Seoul National University) for sharing the primary transcript sequences for let-7c and let-7g. This work was supported by NIH grant R01 GM114147 (to L.J.), NIH R01-GM083300 (to E.C.L.) and MSK Core Grant P30-CA008748. L.J. is an investigator of the Howards Hughes Medical Institute.

## Author contribution

A.G. and L.J. conceived and designed the study. A.G. performed most of the experiments. T.C. assisted A.G. with *in-vitro* qualitative cleavage assays for SRSF3-Drosha interface mutants. R.S. performed the cell-based Northern blot assays under E.C.L. guidance. A.G. and L.J. analyzed the data and wrote the manuscript, and all authors edited the text.

## STAR★METHODS

### KEY RESOURCES TABLE-

| Reagent or Resource | Source | Identifier |
| --- | --- | --- |
| **Chemicals, peptides and recombinant proteins** |  |  |
| [γ-<sup>32</sup>P] ATP | Perkin Elmer | Cat# BLU502Z250UC |
| Strep-Tactin Superflow high-capacity resin | IBA Lifesciences | Cat# 2-1208-025 |
| SYBR Gold Nucleic acid gel stain | ThermoFisher Scientific | Cat# S11494 |
| DSG, No-weight format | ThermoFisher Scientific | Cat# A35392 |
| T4 Polynucleotide Kinase | NEB | Cat# M0201L |
| Proteinase K | NEB | Cat# P8107S |
| RNasin RNase inhibitor | Promega | Cat# N2515 |
| Octyl β-D-glucopyranoside | ThermoFisher Scientific | Cat# BP585-1 |
| HyClone CCM3 media | Cytiva | Cat# SH30065.02 |
| ESF921 cell culture media | Expression systems | Cat# 96-001-01 |
| d-Desthiobiotin | Millipore-Sigma | Cat# D1411-1G |
| HiTrap SP-HP Column | Cytiva | Cat# 17115201 |
| Superose 6 increase 3.2/300 column | Cytiva | Cat# 29091598 |
| Superose 6 increase 10/300 column | Cytiva | Cat# 29091596 |
| Superdex 75 increase 10/300 column | Cytiva | Cat# 29148721 |
| **Deposited data** |  |  |
| Coordinates for MP<sup>he</sup>-pri-let-7a1 structure | This work | PDB: 9ASP |
| Cryo-EM maps for MP<sup>he</sup>-pri-let-7a1 structure | This work | EMD: 43822 |
| Coordinates for MP<sup>he</sup>-pri-miR-98 structure | This work | PDB: 9ASN |
| Cryo-EM maps for MP<sup>he</sup>-pri-miR-98 structure | This work | EMD: 43820 |
| Coordinates for MP<sup>he</sup>-pri-let-7a2 structure | This work | PDB: 9ASO |
| Cryo-EM maps for MP<sup>he</sup>-pri-let-7a2 structure | This work | EMD: 43821 |
| Coordinates for MP<sup>he</sup>-pri-let-7f1 structure | This work | PDB: 9ASM |
| Cryo-EM maps for MP<sup>he</sup>-pri-let-7f1 structure | This work | EMD: 43819 |
| Coordinates for MP<sup>he</sup>-pri-let-7f1-SRSF3 structure | This work | PDB: 9ASQ |
| Cryo-EM maps for MP<sup>he</sup>-pri-let-7f1-SRSF3 structure | This work | EMD: 43823 |
| **Experimental models: Cell lines** |  |  |
| Sf9 insect cells | ThermoFisher Scientific | Cat# 11496015 |
| High Five insect cells | ThermoFisher Scientific | Cat# B85502 |
| *E.coli* DH5α cells (high efficiency) | ThermoFisher Scientific | Cat# 18258-012 |
| *E.coli* DH10Bac cells | ThermoFisher Scientific | Cat# 12033-015 |
| *E.coli* Rosetta 2 (DE3) cells | Novagen | Cat# 71400 |
| **Oligonucleotides** |  |  |
| Pri-miRNA sequences. See Table S2. | N/A | N/A |
| **Recombinant DNA** |  |  |
| pRSF_HH_T7_pri-let-7a1_HDV | This work | N/A |
| pRSF_HH_T7_pri-let-7a2_HDV | This work | N/A |
| pRSF_HH_T7_pri-let-7c_HDV | This work | N/A |
| pRSF_HH_T7_pri-let-7f1_HDV | This work | N/A |
| pRSF_HH_T7_pri-let-7g_HDV | This work | N/A |
| pRSF_HH_T7_pri-miR-98_HDV | This work | N/A |
| pRSF_HH_T7_pri-let-7c<sup>CNNC/UUUU</sup>_HDV | This work | N/A |
| pRSF_HH_T7_pri-let-7g<sup>CNNC/UUUU</sup>_HDV | This work | N/A |
| pRSF_HH_T7_pri-miR-16-1_HDV | This work | N/A |
| pRSF_HH_T7_pri-miR-16-2_HDV | This work | N/A |
| pcDNA4/cmyc-Drosha | Landthaler et al., 2004 | Addgene # 10828 |
| pFL-(Strep)<sub>2</sub>-hDrosha | This work | N/A |
| pFL-(Strep)<sub>2</sub>-hDroshaΔN | This work | N/A |
| pFL-(Strep)<sub>2</sub>-hDroshaΔ<br>N-Mut-Interface1<br>(D758H/E841R) | This work | N/A |
| pFL-(Strep)<sub>2</sub>-hDroshaΔ<br>N-Mut-Interface2<br>(K738E/R923E/Q924A) | This work | N/A |
| pFL-(Strep)<sub>2</sub>-hDroshaΔ<br>N-Mut-Interface1+2<br>(D758H/E841R<br>/K738E/R923E/Q924A) | This work | N/A |
| pFL-SST-hDrosha | This work | N/A |
| pFL-SST-hDroshaΔN | This work | N/A |
| pFLAG/HA-DGCR8 | Landthaler et al., 2004 | Addgene # 10921 |
| pFL-SST-DGRC8 | This work | N/A |
| pSPL-6xHis-DGCR8 | This work | N/A |
| pSPL-6xHis-DGCR8-ΔN1 (175-751) | This work | N/A |
| pSPL-6xHis-DGCR8-ΔN2 (223-751) | This work | N/A |
| pSPL-6xHis-DGCR8-dsRBDs (493-751) | This work | N/A |
| pET28a-TEV-hSRSF3 | Genscript | https://www.genscript.com |
| pET28a-TEV-hSRSF3-RRM (1-84) | This work | N/A |
| pET28a-TEV-hSRSF3-RRM-thumb (1-90) | This work | N/A |
| pET28a-TEV-hSRSF3-Mut1 (E79A/S81A) | This work | N/A |
| pET28a-TEV-hSRSF3-Mut2 (W40A/N44A) | This work | N/A |
| pET28a-TEV-hSRSF3-Mut1+2 (E79A/S81A/W40A/N44A) | This work | N/A |
| pET28a-TEV-hSRSF3-Mut-Interface1 (R69E/R75E) | This work | N/A |
| pET28a-TEV-hSRSF3-Mut-Interface2 (D67H/E79R/R77A) | This work | N/A |
| pET28a-TEV-hSRSF3-Mut-Interface1+2 (R69E/R75E/D67H/E79R/R77A) | This work | N/A |
| **Software and algorithms** |  |  |
| EPU software | ThermoFisher Scientific |  |
| CryoSPARC v2 | Punjani et al., 2017 | https://cryosparc.com |
| WARP | Tegunov and Cramer, 2019 | http://www.warpem.com/warp/ |
| Coot | Emsley and Cowtan, 2004 | https://www2.mrc-lmb.cam.ac.uk/personal/pemsley/coot/ |
| PHENIX 1.20 | Adams et al., 2010 | http://www.phenix-online.org/ |
| ChimeraX | Pettersen et al.,<br>2004 | https://www.cgl.ucsf.edu/chimerax/ |
| PyMol 2.5.5 | Schrödinger LLC | https://pymol.org/2/ |
| MolProbity | Chen et al., 2010. | http://molprobity.biochem.duke.edu/ |
| PRISM 9.5.1 software | GraphPad | https://www.graphpad.com/scientific-software/prism/ |

### Experimental Models and Subject Details

The MuiltiBac baculovirus expression system was used for MP proteins expression in insect cells (Sf9 or HighFive). Baculovirus generated in Sf9 cells was maintained in HyClone CCM3 Cell Culture Media (Cytiva), while HighFive cells were cultured in ESF921 media (Expression systems).

### Expression and purification of human MP-heme variant proteins

Human Drosha isoform 4 and human DGCR8 clones were purchased from Addgene. cDNA encoding different length variants of Drosha were cloned into pFL plasmid and expressed as a N-terminal Dual-strep tag fusion, while DGCR8 cDNA length variants were cloned into pSPL and expressed as N-terminal 6xHis tag fusions. Different combinations of Drosha and DGCR8 clones were Cre-fused and expressed in either Sf9 or HighFive cells using the MultiBac baculovirus expression system. We used N-terminally truncated Drosha^317–1337^ and N-terminally truncated DGCR8^175–751^ for cryo-EM and biochemical studies, as these truncations does not impact the *in-vitro* catalytic activity of MP^16^ and has been used in previous studies^23,54^. The insect cells were infected with baculovirus at 27°C for 60 hrs, and supplemented with 0.75 mM 5-aminoleuvelinic acid (5-ALA), an intermediated in heme-biosynthesis pathway, to enrich the MP protein with heme (MP^he^) during expression. Insect cells were harvested in resuspension buffer (50 mM Tris pH 8.0, 100 mM NaCl and 5 mM DTT) supplemented with protease inhibitor (PI) mix (Pepstatin, Leupeptin, PMSF, Benzamidine and Aprotinin) before flash freezing in liquid N2. The cells were thawed, 650 mM NaCl and 10% glycerol were added before sonication. After ultracentrifugation at 40,000 rpm for 1 hr, cleared lysate was loaded on 4 ml Strep-Tactin Superflow beads (IBA lifesciences) and washed extensively before co-eluting the Drosha and DGCR8 (MP) in 25 mM HEPES pH 7.5, 200 mM NaCl, 5 mM DTT, 10% glycerol supplemented with 7 mM desthiobiotin. The eluted MP protein was diluted with an equal volume of dilution buffer (50 mM Bis-Tris pH 6.8, 5 mM DTT and 10% glycerol) and loaded onto the HiTrap SP HP cation exchange column (Cytiva) pre-equilibrated in Buffer-A (25 mM Tris pH 6.8, 75 mM NaCl, 5 mM DTT and 10% glycerol). A linear NaCl gradient (from 75 mM to 1 M) was used to elute the MP.

Low-heme MP (MP^apo^) and heme-bound MP (MP^hb^) had different ionic strengths and were separated during HiTrap SP-HP column chromatography, as observed from Abs450 peak and A450/280 ratios. The heme-enriched MP (MP^he^) protein eluted as a single peak with a higher Abs450 and A450/280 ratio. Eluted heme-variant MP proteins were separated, analyzed on SDS-PAGE, pooled and loaded onto Superose 6 increase 10/300 gel filtration column (Cytiva) pre-equilibrated with SEC buffer (25 mM HEPES pH 7.5, 400 mM NaCl, 5 mM DTT and 10% glycerol). Peak fractions for eluted MP proteins were concentrated and flash frozen in liquid N_2_ and stored at -80°C

### Expression and purification of SRSF3 proteins

The human SRSF3-pET28a clone was purchased from Genscript (Piscataway, NJ). Different SRSF3 variants were subcloned in pET28a and expressed as TEV protease-cleavable N-terminal 6xHis-tag fusions in *E.coli* Rosetta2(DE3) cells (Novagen). *E.coli* cells were grown in TB media up to an OD_600_ of ∼ 1.2 at 37°C, and protein expression was induced for 16 hr at 18°C by adding 0.5 mM IPTG. Cells were harvested in *E.coli* resuspension buffer (25 mM Tris pH 8.0, 500 mM NaCl, 5 mM ß-Me and 10 mM Imidazole), supplemented with PI cocktail (pepstatin, leupeptin, PMSF, benzamidine and aprotinin) before flash freezing in liquid N_2_. The cells were thawed and lysed by sonication and bulk nucleic acids were precipitated by mixing 0.2% PEI (Poly-ethylenimine) into the lysate before ultracentrifugation at 40,000 rpm for 1 hr. Using the cleared lysate, Ni-NTA affinity chromatography was performed to elute 6xHis-SRSF3 protein in Ni-NTA elution buffer (25 mM HEPES pH 7.5, 200 mM NaCl, 5 mM ß-Me, 250 mM imidazole and 5% glycerol). The eluted protein was diluted to reduce the NaCl concentration to 50 mM using dilution buffer (25 mM HEPES pH 7.4, 5 mM ß-Me and 5% glycerol) and loaded onto HiTrap SP-HP column (Cytiva) pre-equilibrated with cation exchange buffer-A (25 mM HEPES pH 7.4, 50 mM NaCl, 5 mM ß-Me and 5% glycerol). NaCl linear gradient from 50 mM – 1000 mM was used to elute the 6xHis-SRSF3 protein and analyzed on SDS-PAGE. Fractions containing pure protein were pooled, concentrated and injected into the Superdex 75 increase 10/300 column (Cytiva) pre-equilibrated with SEC buffer (25 mM HEPES pH 7.5, 150 mM NaCl and 1 mM DTT). Purified SRSF3 protein was concentrated before flash freezing in liquid N_2_.

### MicroKit SEC for MP heme-variants

Different MP heme-variants were analyzed by injecting 2.4 μM (12 μg total protein) of purified protein into Superose 6 increase 3.2/300 column (Cytiva) pre-equilibrated in Microkit SEC buffer (HEPES 25 mM pH 7.5, NaCl 400 mM, DTT 5 mM, 10% glycerol and 10 μM ZnCl_2_). Absorption at UV280, UV260 and UV450 were recorded, while UV280 and UV450 traces were used to plot the final chromatograms. The A450/280 ratio was calculated from the values at the tip of each chromatogram peak. N=3.

### In-vitro transcription (IVT) of pri-miRNAs

The cDNAs encoding different pri-let-7 miRNAs were subcloned in pRSF plasmid, with a flanking 5’ hammerhead (HH) ribozyme and 3’ hepatitis delta virus (HDV) ribozyme sequences. Large scale *in-vitro* transcription reactions were run using in-house purified T7 RNA polymerase. After transcription, DNA templates were digested using RNase-free DNase (NEB), and RNA was phenol-chloroform extracted followed by isopropanol precipitation (overnight at -80°C). The RNA pellet was dissolved in TE buffer, and PAGE purified from a 6% denaturing polyacrylamide gel in RNase-free water. Purified pri-miRNA was concentrated using Amicon Ultra-14 (10 kDa MWCO) and stored at -80°C.

### Qualitative cleavage assays for pri-miRNAs

*In-vitro* pri-miRNA time course cleavage assays were performed by incubating ∼150 nM pri-miRNA with ∼300 nM purified MP protein in processing assay buffer (20 mM HEPES pH 7.5, 100 mM NaCl, 2.5 mM DTT, 5 mM MgCl2, 2 U/ml RNasin and 10% glycerol) at a final volume of 100 μl. The reaction mix was incubated at 37°C and 15 μl aliquots were removed at indicated time points, quenched into stop buffer (1.8% SDS, 10 mM EDTA) followed by Proteinase-K (2 units total) (NEB) treatment for 45 min at 50°C. The samples were diluted with the 2X denaturing buffer (80 % formamide, 1.5 M urea, 2 mM EDTA, 0.05% bromophenol-blue and 0.05% xylene-cyanole) and analyzed on a 12% urea-PAGE (20 cm X 20 cm) (National diagnostics). Gels were stained with SybrGold stain and visualized on BioRad ChemiDoc imaging system.

For qualitative RNA cleavage assays with SRSF3, ∼150 nM pri-miRNA was first incubated with 350 nM SRSF3 protein for 15 min at 30°C in processing assay buffer. Catalysis was initiated by adding ∼250 nM MP protein, and 15 μl aliquots were removed at indicated time points for analysis.

### Quantitative biochemical analysis for pri-let-7 processing

The quantitative biochemical analyses for different pri-miRNAs were done in near pre-steady state conditions to study single turnover kinetics for M^2^P^2^. For all kinetics assays, 135 μl volume reaction was set up by incubating 50 nM of annealed pri-miRNA substrate in processing assay buffer (20 mM HEPES pH 7.5, 100 mM NaCl, 2.5 mM DTT, 5 mM MgCl_2_, 1 U/ml RNasin and 10% glycerol) at 30°C for 5 mins. The reaction was initiated by adding 500 nM of MP^he^, and 15 μl aliquots were removed at 0.25’, 0.5’, 1’, 2’, 5’, 10’, 20’, 45’ and quenched into 3 ***μ***l of 5X stop buffer (9% SDS and 50 mM EDTA). For assays involving the MP^apo^, the samples were collected at 0.25’, 0.5’, 1’, 2’, 5’, 10’, 30’, 60’. The samples were further treated with 2U Proteinase-K (NEB) for 45 min at 50°C before diluting the samples with 2X denaturing buffer (80 % formamide, 1.5 M urea, 2 mM EDTA, 0.05% bromophenol-blue and 0.05% xylene-cyanole). Samples were heated at 95°C for 5 mins before analyzing onto the 12% denaturing urea-PAGE (National diagnostics).

For the kinetics assays testing the effect of SRSF3 on M^2^P^2^, we pre-incubated increasing concentrations of SRSF3-FL protein with 50 nM pri-miRNA in 1:2.5, 1:12.5 and 1:25 molar ratio (125 nM, 625 nM and 1250 nM of SRSF3-FL respectively) for 10 mins at 30°C, before starting the reaction by adding 500 nM MP^he^. 15 μl aliquots were removed at 0.25’, 0.5’, 1’, 2’, 5’, 10’, 20’, 45’ and quenched into 3 μl of 5X stop buffer, processed and analyzed as mentioned above.

For testing the effect of different mutants and truncations in SRSF3 on M^2^P^2^, 625 nM of SRSF3 protein was pre-incubated with 50 nM pri-miRNA substrate for 10 mins at 30°C, before starting the reaction by adding 500 nM MP^he^. 15 μl aliquots were removed at 0.25’, 0.5’, 1’, 2’, 5’, 10’, 20’, 45’ and quenched into 3 μl of 5X stop buffer, processed and analyzed as mentioned above.

Gels were stained with SybrGold and visualized on a BioRad ChemiDoc imaging system. All the RNA products formed after the reactions were quantified in each lane in the gels by ImageJ software, and the relative fraction of leftover pri-miRNA substrate, generated pre-miRNA (productive cleavage), and the unproductive cleavage product were calculated. GraphPad Prism 9 was used for data and statistical analysis. Quantified data were fitted to a mono-phasic decay equation, to calculate the lower limit and upper limit values at 95% confidence interval (CI), from which a best-fit observed rate constant (k_obs_) is determined in GraphPad Prism 9. The produced pre-let-7c product in different conditions was further cleaved by MP overtime, so the traces were not fitted to any decay equation and are shown only with connecting lines in Figure 5D and 6G.

For testing the Q1144A mutation’s effect on M^2^P^2^, the 1 nM of 5’ ^32^P-labelled pri-let-7a1 was mixed with 10 nM of MP^he^ in 100 μl volume, and 10 μl aliquots were collected at 0.16’, 0.5’, 1’, 2’, 5’, 10’, 15’, 30’ and 60’ and directly quenched into stop buffer. Samples were processed as mentioned above, analyzed onto the 12% denaturing urea-PAGE (National diagnostics) and imaged via phosphor-imaging on Typhoon FLA 7000 (GE healthcare Life Sciences)

### Electrophoretic Mobility Shift assays (EMSA)

For EMSA, ∼150 nM pri-let-7 RNA was mixed with different MP^he^ proteins in a 1:1, 1:2 and 1:4 molar ratios in EMSA buffer-1 (25 mM HEPES pH 7.5, 100 mM NaCl, 2.5 mM DTT, 5 mM CaCl_2_, 10 % glycerol, 0.01 % Triton X-100 and 1 U/ml RNasin). The reaction was incubated on ice for 1 hr, 20 % glycerol was added and analyzed on a 5 % TBE gel (Bio-Rad) run in ice-cold 0.5 X TBE buffer. Gels were stained with SybrGold and visualized on BioRad ChemiDoc imaging system.

For EMSA with SRSF3, 1 nM of the 5’ ^32^P-labelled pri-miRNA was incubated with SRSF3 proteins at the indicated concentrations in EMSA buffer (25 mM HEPES pH 7.5, 75 mM NaCl, 2 mM DTT, 8% glycerol, 0.01 % Triton X-100, 1 mM EDTA, 0.1 mg/ml BSA) for 1 hr at 4°C. EMSA loading dye (50 % glycerol, 0.05 % xylene cyanol and 0.05 % bromophenol blue) was added and samples analyzed on a 20x20 cm 4 % TBE gel run in ice-cold 0.5 X TBE buffer. Gels were exposed to phosphor screens overnight and imaged using Typhoon FLA 7000 (GE healthcare Life Sciences).

### In-vitro pull-down assays

Pull-down assays were performed with MP^he^, pri-let-7f1 against different SRSF3 proteins (WT or mutants as marked in the figures). 25 μl Strep-Tactin beads were coated with BSA (0.2 mg/ml) for 20 mins, followed by incubation with 4 μM MP^he^ for 15 mins on ice. The mixture was centrifuged at 200 xg for 4 min, and excess solution (FT-MP^he^+BSA) was removed. Beads were then incubated with 20 μM of different SRSF3 protein with or without 6 μM of pri-let-7f1 for 20 mins on ice. The flow-through (FT-SRSF3-f1) was collected before sequentially washing the beads with 200 μl, 400 μl & 500 μl wash buffer (25 mM HEPES pH 7.5, 200 mM NaCl, 2 mM DTT, 5 mM CaCl_2_, 10% glycerol, 0.01% Triton X-100). Bound complexes were eluted twice in 30 μl elution buffer (25 mM HEPES pH 7.5, 100 mM NaCl, 2 mM DTT, 5 mM CaCl_2_, 10% glycerol, 0.01% Triton X-100 and 5 mM desthio-biotin) and analyzed on SDS-PAGE and urea-PAGE.

### Northern blots

Co-transfection of the Drosha-KO HEK293T cells^55^ with miRNA plasmids (for each well: 300 ng of WT or CNNC mutant miRNAs with 50 ng of non-CNNC mir-24 plasmids) and Drosha plasmids (800 ng per well) was performed in 12-well cell culture plates using Lipofectamine2000 (Thermo Fisher) according to the manufacturer’s protocol. Cells were harvested 48 hours post-transfection using Trizol reagent for total RNA extraction. Equal amounts of total RNAs (10 µg) were denatured at 95°C and fractionated by electrophoresis on a 20% urea polyacrylamide gel in 0.5x TBE buffer. The gel was transferred to GeneScreen Plus membrane (Perkin Elmer) at 300 mA for 1.5 hr, UV-crosslinked and baked at 80°C for 30 min and hybridized with γ-^32^P-labeled miRNA probes at 42°C overnight (hybridization buffer: 5x SSC, 7% SDS, 2x Denhardt’s solution). The membrane was washed with Non-Stringent Wash Solution (3x SSC, 5% SDS, 10x Denhardt’s solution) followed by two rounds of wash with Stringent Wash Solution (1x SSC, 1% SDS). Each wash step was conducted at 42°C for 30 min. The membrane was sealed in plastic wrap, inserted into a film cassette and exposed for 1∼3 days. For re-probing of the same blot with other miRNA probes, the blot was washed with 1% SDS at 80°C for 30 min and then hybridized with the following miRNA probes: anti-mir-16-5p CGCCAATATTTACGTGCTGCTA; anti-let-7g-5p AACTGTACAAACTACTACCTCA; anti-mir-24-3p CTGTTCCTGCTGAACTGAGCCA.

### Cryo-EM sample preparation

Pri-let-7s were annealed by heating at 85°C for 5 min and snap cooling on ice for 10 mins. MP^he^ and different pri-let-7s were mixed in 1:1.25 molar ratios in cryo-EM reaction buffer (25 mM HEPES pH 7.5, 100 mM NaCl, 5 mM DTT, 5 mM CaCl_2_) and incubated for 1:30 hr on ice. The complex was crosslinked using 2 mM DSG (ThermoFisher Scientific) for 25 mins at 20°C. The crosslinked complex was separated from excess RNA via SEC over Superose 6 increase 10/300 column (Cytiva) pre-equilibrated in cryo-EM sample buffer (20 mM HEPES pH 7.5, 120 mM NaCl, 5 mM DTT, 5 mM CaCl_2_) and concentrated to ∼1.0 – 1.5 mg/ml (Bradford method) before grid preparation. For MP^he^-pri-let-7f1-SRSF3 sample, 1.4 fold molar excess SRSF3-FL to RNA was added while assembling the complex before crosslinking. For MP^he^-pri-let-7a2 sample, SRSF3-RRM was added in the sample, but no density was observed for the SRSF3-RRM domain.

For cryo-EM grid preparation, 0.05% w/v β-OG (Octyl β-D-glucopyranoside) (ThermoFisher Scientific) detergent were added prior to application of 3 μl sample onto a glow-discharged Quantifoil R 0.6/1 300 mesh grid (for MP^he^-pri-let-7a1, MP^he^-pri-let-7f1, MP^he^-pri-let-7f1-SRSF3) or glow-discharged UltrAuFoil R 0.6/1 300 mesh grid (for MP^he^-pri-miR-98, MP^he^-pri-let-7a2, MP^he^-pri-let-7f1-SRSF3), incubated for 10 sec at 25°C and 95% humidity, blotted for 3.1 s, and plunged into liquid ethane using an Automatic Plunge Freezer EM GP2 (Leica).

### Cryo-EM data acquisition

Cryo-EM data were collected on a Titan Krios transmission electron microscope (ThermoFisher Scientific) operating at 300 keV. EPU data collection software (v 2.10.0.5) (ThermoFisher Scientific) was used, and dose-fractionated movies were collected using a K3 direct electron detector (Gatan) operating in electron counting mode.

For MP^he^-pri-let-7a1, 30-framed movies were collected with an exposure rate of 2.405 e^−^/Å^2^/frame resulting in a cumulative exposure of 72.16 e^−^/Å^2^. A total 11,160 micrographs were collected at 81,000x magnification (1.1 Å/pixel) and defocus range of 0.6 to 2.2 μm.

For MP^he^-pri-miR-98, 30-framed movies were collected with an exposure rate of 1.42 e^−^/Å^2^/frame resulting in a cumulative exposure of 42.6 e^−^/Å^2^. A total 11,051 micrographs were collected at 81,000x magnification (1.1 Å/pixel) and defocus range of 0.7 to 2.2 μm. For MP^he^-pri-let-7a2, 30-framed movies were collected with an exposure rate of 2.6 e^−^/Å^2^/frame resulting in a cumulative exposure of 78 e^−^/Å^2^. A total 5,486 micrographs were collected at 105,000x magnification (0.856 Å/pixel) and defocus range of 0.6 to 2.0 μm.

For MP^he^-pri-let-7f1, 30-framed movies were collected with an exposure rate of 2.59 e^−^/Å^2^/frame resulting in a cumulative exposure of 77.6 e^−^/Å^2^. A total 8,217 micrographs were collected at 105,000x magnification (0.856 Å/pixel) and defocus range of 0.6 to 2.2 μm in CDS (correlative double sampling) mode.

For MP^he^-pri-let-7f1-SRSF3, 30-framed movies were collected with an exposure rate of 2.55 e^−^/Å^2^/frame resulting in a cumulative exposure of 76.6 e^−^/Å^2^. Total 7,995 and 6,429 micrographs were collected at 105,000x magnification (0.856 Å/pixel) and defocus range of 0.7 to 2.2 μm from a QuantiFoil and UltrAuFoil grid respectively.

### Cryo-EM Image processing

WARP (v 1.0.9) was used for real-time image pre-processing (motion correction, CTF estimation, and particle picking)^56^ for all the MP^he^-pri-let-7 miRNA structures. Particle picking was performed with the BoxNet pretrained neural network bundle included in WARP that is implemented in TensorFlow. A particle diameter of 180 Å and threshold score of 0.6 yielded 1,749,636 particle coordinates for MP^he^-pri-let-7a1, 967,368 particle coordinates for MP^he^-pri-miR-98, 1,742,639 particle coordinates for MP^he^-pri-let-7f1, 771,705 particle coordinates for MP^he^-pri-let-7a2. For MP^he^-pri-let-7f1-SRSF3, WARP processing yielded 846,678 and 871,735 particle coordinates for the QuantiFoil and UltrAufoil datasets respectively. All subsequent processing steps were carried out in cryoSPARC v3.2^57^.

For all the structures, extracted particles were 2D classified, and a subset of those were used for ab-initio 3D reconstruction after manually inspecting each 2D class. Separating particles into total a 8-10 ab-initio classes was critical in improving the map quality for all the structures in this study. The resulting models were then used for 3D heterogeneous refinement against the whole particle set. Thus, we separated 397,113 particles for MP^he^-pri-let-7a1 (Figure S2D), 209,910 particles for MP^he^-pri-miR-98 (Figure S2A), 459,947 particles for MP^he^-pri-let-7f1 (Figure S2G) and 251,413 particles for MP^he^-pri-let-7a2 (Figure S4A) for further refinements. For MP^he^-pri-let-7f1-SRSF3 3D classification yielded 254,462 and 234,705 particles from the QuantiFoil and UltrAuFoil datasets respectively (Figure S6A).

For MP^he^-pri-let-7a1, 397,113 particles were refined with 4 random decoy classes, followed by a homogeneous and a non-uniform refinement yielding a cryo-EM map of 3.2 Å resolution according to gold standard FSC (GSFSC) (Figure S2E).

For MP^he^-pri-miR-98 and MP^he^-pri-let-7f1, 3D classified particle stacks (209,910 and 459,947 particles respectively) were iteratively processed by 2 cycles of heterogeneous and homogeneous refinement separating 173,800 and 378,162 particles for their respective structures. This particle stack was non-uniformly refined to generate 3.2 Å and 2.9 Å resolution cryo-EM maps (according to GSFSC) for MP^he^-pri-miR-98 (Figure S2B) and MP^he^-pri-let-7f1 (Figure S2H) respectively.

Similarly, for MP^he^-pri-let-7a2, one homogeneous and non-uniform refinement of 251,413 particles yielded the final 2.9 Å cryo-EM map according to GSFSC (Figure S4A-B). For MP^he^-pri-let-7f1-SRSF3, the two 3D refined particle stacks were merged (total 489,167 particles) and taken for 2 cycles of iterative heterogeneous and homogeneous refinements separating 325,534 good particles. Finally, a non-uniform refinement was performed to generated cryo-EM map of 3.1 Å resolution, according to GSFSC (Figure S6).

We further performed local sharpening and de-noising of cryo-EM maps using non-linear post-processing with <u>Deep</u> cryo-<u>EM</u> Map En<u>hancer</u> (DeepEMhancer)^58^. These DeepEMhanc’ed maps had much improved cryo-EM density for Drosha, DGCR8, and the apical loop of the RNA (Figure S2C, S2F, S2I). Sharpened maps were used for model visualizations and building, while original maps were used for structure model refinements.

### Model-building, refinement and validation

The atomic model of MP-pri-miR-16-2 (PDBid 6V5B)^23^ was used as a starting model for the MP^he^-pri-miR-98 structure, which was rigid body fitted in the map using ChimeraX^59^. Although Drosha (from MP-pri-miR-16-2) aligned well in the cryo-EM map, several regions had to be rebuilt to get the final MP^he^-pri-miR-98 structure model (Figure 2C), which was then used as template for the other MP^he^-let-7 structures in this study. Though the nucleotide register was easily identifiable in the maps, we also utilized the M-fold server^60^ as a guide for building a few base-pairs in RNA upper stem only. The atomic model of the let-7 pre-miRNA apical loop from the Lin28-pre-let-7 crystal structure (PDBid 5UDZ)^39^ was fitted into the MP^he^-pri-let-7a1 map using ChimeraX, and individual nucleotides were replaced to match the appropriate sequence in Coot (v 0.9.4)^61^. For the MP^he^-pri-let-7f1-SRSF3 structure the extended 3p RNA fragment was manually build using Coot. The atomic model of the SRSF3 RRM domain (PDBid 2I2Y)^42^ was used to rigid body fit into the map density near the Drosha PAZ-like domain using ChimeraX and manually build further. All atomic model building was done in Coot, and refinements were performed in PHENIX (v1.20.1-4487-000)^62,63^. Secondary structure restraints for protein and RNA were used throughout the refinement process. The DeepEMhance’d maps were also utilized for visualizing and building the structure models, while we used the unsharpened cryo-EM map for refinements. Structure validation was done using the MolProbity server^64^. The structure figures were generated by ChimeraX and PyMOL molecular graphics system (Version 2.5.5, Schrödinger, LLC, Heidelberg, D). The data collection and model statistics are summarized in Table S1.

### Lead contact and Material availability

Further information and requests for resources and reagents should be directed to and will be fulfilled by the Lead Contact, Leemor Joshua-Tor. Reagents generated in this study are available from the Lead Contact with a completed Materials Transfer Agreement.

### Limitation of the study

Though we can resolve more of the HBD of Drosha that was reported previously, a significant portion of that domain is still untraceable. In addition, we could not identify the heme itself. Therefore, the interplay between heme and the UGUG motif, in particular, needs further characterization. Though a good representation of let-7 pri-miRNAs were used in this study, other pri-miRNAs may add to a more complete understanding of how the MP recognizes pri-miRNAs and how that may affect M^2^P^2^.

## References

1 Shang, R., Lee, S., Senavirathne, G. & Lai, E. C. microRNAs in action: biogenesis, function and regulation. Nat Rev Genet 24, 816–833, doi:10.1038/s41576-023-00611-y (2023).

2 Gregory, R. I. et al. The Microprocessor complex mediates the genesis of microRNAs. Nature 432, 235–240, doi:10.1038/nature03120 (2004).

3 Lee, Y. et al. The nuclear RNase III Drosha initiates microRNA processing. Nature 425, 415–419, doi:10.1038/nature01957 (2003).

4 Denli, A. M., Tops, B. B., Plasterk, R. H., Ketting, R. F. & Hannon, G. J. Processing of primary microRNAs by the Microprocessor complex. Nature 432, 231–235, doi:10.1038/nature03049 (2004).

5 Okada, C. et al. A high-resolution structure of the pre-microRNA nuclear export machinery. Science 326, 1275–1279, doi:10.1126/science.1178705 (2009).

6 Lee, Y. Y., Lee, H., Kim, H., Kim, V. N. & Roh, S. H. Structure of the human DICER-pre-miRNA complex in a dicing state. Nature 615, 331–338, doi:10.1038/s41586-023-05723-3 (2023).

7 Czech, B. et al. Hierarchical rules for Argonaute loading in Drosophila. Mol Cell 36, 445–456, doi:10.1016/j.molcel.2009.09.028 (2009).

8 Bartel, D. P. MicroRNAs: target recognition and regulatory functions. Cell 136, 215–233, doi:10.1016/j.cell.2009.01.002 (2009).

9 Lai, E. C. Micro RNAs are complementary to 3’ UTR sequence motifs that mediate negative post-transcriptional regulation. Nat Genet 30, 363–364, doi:10.1038/ng865 (2002).

10 Zeng, Y. & Cullen, B. R. Efficient processing of primary microRNA hairpins by Drosha requires flanking nonstructured RNA sequences. J Biol Chem 280, 27595–27603, doi:10.1074/jbc.M504714200 (2005).

11 Zeng, Y., Yi, R. & Cullen, B. R. Recognition and cleavage of primary microRNA precursors by the nuclear processing enzyme Drosha. EMBO J 24, 138–148, doi:10.1038/sj.emboj.7600491 (2005).

12 Han, J. et al. Molecular basis for the recognition of primary microRNAs by the Drosha-DGCR8 complex. Cell 125, 887–901, doi:10.1016/j.cell.2006.03.043 (2006).

13 Fang, W. & Bartel, D. P. The Menu of Features that Define Primary MicroRNAs and Enable De Novo Design of MicroRNA Genes. Mol Cell 60, 131–145, doi:10.1016/j.molcel.2015.08.015 (2015).

14 Ma, H., Wu, Y., Choi, J. G. & Wu, H. Lower and upper stem-single-stranded RNA junctions together determine the Drosha cleavage site. Proc Natl Acad Sci U S A 110, 20687–20692, doi:10.1073/pnas.1311639110 (2013).

15 Auyeung, V. C., Ulitsky, I., McGeary, S. E. & Bartel, D. P. Beyond secondary structure: primary-sequence determinants license pri-miRNA hairpins for processing. Cell 152, 844–858, doi:10.1016/j.cell.2013.01.031 (2013).

16 Nguyen, T. A. et al. Functional Anatomy of the Human Microprocessor. Cell 161, 1374–1387, doi:10.1016/j.cell.2015.05.010 (2015).

17 Kwon, S. C. et al. Molecular Basis for the Single-Nucleotide Precision of Primary microRNA Processing. Mol Cell 73, 505–518 e505, doi:10.1016/j.molcel.2018.11.005 (2019).

18 Kim, K. et al. A quantitative map of human primary microRNA processing sites. Mol Cell 81, 3422–3439 e3411, doi:10.1016/j.molcel.2021.07.002 (2021).

19 Kwon, S. C. et al. Structure of Human DROSHA. Cell 164, 81–90, doi:10.1016/j.cell.2015.12.019 (2016).

20 Kim, K., Nguyen, T. D., Li, S. & Nguyen, T. A. SRSF3 recruits DROSHA to the basal junction of primary microRNAs. RNA 24, 892–898, doi:10.1261/rna.065862.118 (2018).

21 Lee, H., Han, S., Kwon, C. S. & Lee, D. Biogenesis and regulation of the let-7 miRNAs and their functional implications. Protein Cell 7, 100–113, doi:10.1007/s13238-015-0212-y (2016).

22 Jin, W., Wang, J., Liu, C. P., Wang, H. W. & Xu, R. M. Structural Basis for pri-miRNA Recognition by Drosha. Mol Cell 78, 423–433 e425, doi:10.1016/j.molcel.2020.02.024 (2020).

23 Partin, A. C. et al. Cryo-EM Structures of Human Drosha and DGCR8 in Complex with Primary MicroRNA. Mol Cell 78, 411–422 e414, doi:10.1016/j.molcel.2020.02.016 (2020).

24 Le, M. N., Nguyen, T. D. & Nguyen, T. A. SRSF7 and SRSF3 depend on RNA sequencing motifs and secondary structures to regulate Microprocessor. Life Sci Alliance 6, doi:10.26508/lsa.202201779 (2023).

25 Faller, M., Matsunaga, M., Yin, S., Loo, J. A. & Guo, F. Heme is involved in microRNA processing. Nat Struct Mol Biol 14, 23–29, doi:10.1038/nsmb1182 (2007).

26 Weitz, S. H., Gong, M., Barr, I., Weiss, S. & Guo, F. Processing of microRNA primary transcripts requires heme in mammalian cells. Proc Natl Acad Sci U S A 111, 1861–1866, doi:10.1073/pnas.1309915111 (2014).

27 Barr, I. et al. Ferric, not ferrous, heme activates RNA-binding protein DGCR8 for primary microRNA processing. Proc Natl Acad Sci U S A 109, 1919–1924, doi:10.1073/pnas.1114514109 (2012).

28 Senturia, R. et al. Structure of the dimerization domain of DiGeorge critical region 8. Protein Sci 19, 1354–1365, doi:10.1002/pro.414 (2010).

29 Senturia, R., Laganowsky, A., Barr, I., Scheidemantle, B. D. & Guo, F. Dimerization and heme binding are conserved in amphibian and starfish homologues of the microRNA processing protein DGCR8. PLoS One 7, e39688, doi:10.1371/journal.pone.0039688 (2012).

30 Partin, A. C. et al. Heme enables proper positioning of Drosha and DGCR8 on primary microRNAs. Nat Commun 8, 1737, doi:10.1038/s41467-017-01713-y (2017).

31 Bussing, I., Slack, F. J. & Grosshans, H. let-7 microRNAs in development, stem cells and cancer. Trends Mol Med 14, 400–409, doi:10.1016/j.molmed.2008.07.001 (2008).

32 Heo, I. et al. Mono-uridylation of pre-microRNA as a key step in the biogenesis of group II let-7 microRNAs. Cell 151, 521–532, doi:10.1016/j.cell.2012.09.022 (2012).

33 Faehnle, C. R., Walleshauser, J. & Joshua-Tor, L. Multi-domain utilization by TUT4 and TUT7 in control of let-7 biogenesis. Nat Struct Mol Biol 24, 658–665, doi:10.1038/nsmb.3428 (2017).

34 Thornton, J. E. et al. Selective microRNA uridylation by Zcchc6 (TUT7) and Zcchc11 (TUT4). Nucleic Acids Res 42, 11777–11791, doi:10.1093/nar/gku805 (2014).

35 Park, J. E. et al. Dicer recognizes the 5’ end of RNA for efficient and accurate processing. Nature 475, 201–205, doi:10.1038/nature10198 (2011).

36 Jumper, J. et al. Highly accurate protein structure prediction with AlphaFold. Nature 596, 583–589, doi:10.1038/s41586-021-03819-2 (2021).

37 Abramson, J. et al. Accurate structure prediction of biomolecular interactions with AlphaFold 3. Nature 630, 493–500, doi:10.1038/s41586-024-07487-w (2024).

38 Fang, W. & Bartel, D. P. MicroRNA Clustering Assists Processing of Suboptimal MicroRNA Hairpins through the Action of the ERH Protein. Mol Cell 78, 289–302 e286, doi:10.1016/j.molcel.2020.01.026 (2020).

39 Wang, L. et al. LIN28 Zinc Knuckle Domain Is Required and Sufficient to Induce let-7 Oligouridylation. Cell Rep 18, 2664–2675, doi:10.1016/j.celrep.2017.02.044 (2017).

40 Mayr, F., Schutz, A., Doge, N. & Heinemann, U. The Lin28 cold-shock domain remodels pre-let-7 microRNA. Nucleic Acids Res 40, 7492–7506, doi:10.1093/nar/gks355 (2012).

41 Nam, Y., Chen, C., Gregory, R. I., Chou, J. J. & Sliz, P. Molecular basis for interaction of let-7 microRNAs with Lin28. Cell 147, 1080–1091, doi:10.1016/j.cell.2011.10.020 (2011).

42 Hargous, Y. et al. Molecular basis of RNA recognition and TAP binding by the SR proteins SRp20 and 9G8. EMBO J 25, 5126–5137, doi:10.1038/sj.emboj.7601385 (2006).

43 Maris, C., Dominguez, C. & Allain, F. H. The RNA recognition motif, a plastic RNA-binding platform to regulate post-transcriptional gene expression. FEBS J 272, 2118–2131, doi:10.1111/j.1742-4658.2005.04653.x (2005).

44 Chen, Y. et al. SYNCRIP, a new player in pri-let-7a processing. RNA 26, 290–305, doi:10.1261/rna.072959.119 (2020).

45 Kawahara, H. et al. Musashi1 cooperates in abnormal cell lineage protein 28 (Lin28)-mediated let-7 family microRNA biogenesis in early neural differentiation. J Biol Chem 286, 16121–16130, doi:10.1074/jbc.M110.199166 (2011).

46 Viswanathan, S. R., Daley, G. Q. & Gregory, R. I. Selective blockade of microRNA processing by Lin28. Science 320, 97–100, doi:10.1126/science.1154040 (2008).

47 Nikolov, D. B. et al. Crystal structure of a TFIIB-TBP-TATA-element ternary complex. Nature 377, 119–128, doi:10.1038/377119a0 (1995).

48 Feng, X. et al. The structure of ORC-Cdc6 on an origin DNA reveals the mechanism of ORC activation by the replication initiator Cdc6. Nat Commun 12, 3883, doi:10.1038/s41467-021-24199-1 (2021).

49 Manley, J. L. & Krainer, A. R. A rational nomenclature for serine/arginine-rich protein splicing factors (SR proteins). Genes Dev 24, 1073–1074, doi:10.1101/gad.1934910 (2010).

50 Zhou, Z. et al. Emerging Roles of SRSF3 as a Therapeutic Target for Cancer. Front Oncol 10, 577636, doi:10.3389/fonc.2020.577636 (2020).

51 Kielkopf, C. L., Lucke, S. & Green, M. R. U2AF homology motifs: protein recognition in the RRM world. Genes Dev 18, 1513–1526, doi:10.1101/gad.1206204 (2004).

52 Kielkopf, C. L., Rodionova, N. A., Green, M. R. & Burley, S. K. A novel peptide recognition mode revealed by the X-ray structure of a core U2AF35/U2AF65 heterodimer. Cell 106, 595–605, doi:10.1016/s0092-8674(01)00480-9 (2001).

53 Selenko, P. et al. Structural basis for the molecular recognition between human splicing factors U2AF65 and SF1/mBBP. Mol Cell 11, 965–976, doi:10.1016/s1097-2765(03)00115-1 (2003).

54 Rice, G. M., Shivashankar, V., Ma, E. J., Baryza, J. L. & Nutiu, R. Functional Atlas of Primary miRNA Maturation by the Microprocessor. Mol Cell 80, 892–902 e894, doi:10.1016/j.molcel.2020.10.028 (2020).

55 Shang, R. & Lai, E. C. Parameters of clustered suboptimal miRNA biogenesis. Proc Natl Acad Sci U S A 120, e2306727120, doi:10.1073/pnas.2306727120 (2023).

56 Tegunov, D. & Cramer, P. Real-time cryo-electron microscopy data preprocessing with Warp. Nat Methods 16, 1146–1152, doi:10.1038/s41592-019-0580-y (2019).

57 Punjani, A., Rubinstein, J. L., Fleet, D. J. & Brubaker, M. A. cryoSPARC: algorithms for rapid unsupervised cryo-EM structure determination. Nat Methods 14, 290–296, doi:10.1038/nmeth.4169 (2017).

58 Sanchez-Garcia, R. et al. DeepEMhancer: a deep learning solution for cryo-EM volume post-processing. Commun Biol 4, 874, doi:10.1038/s42003-021-02399-1 (2021).

59 Pettersen, E. F. et al. UCSF ChimeraX: Structure visualization for researchers, educators, and developers. Protein Sci 30, 70–82, doi:10.1002/pro.3943 (2021).

60 Zuker, M. Mfold web server for nucleic acid folding and hybridization prediction. Nucleic Acids Res 31, 3406–3415, doi:10.1093/nar/gkg595 (2003).

61 Emsley, P. & Cowtan, K. Coot: model-building tools for molecular graphics. Acta Crystallogr D Biol Crystallogr 60, 2126–2132, doi:10.1107/S0907444904019158 (2004).

62 Liebschner, D. et al. Macromolecular structure determination using X-rays, neutrons and electrons: recent developments in Phenix. Acta Crystallogr D Struct Biol 75, 861–877, doi:10.1107/S2059798319011471 (2019).

63 Adams, P. D. et al. PHENIX: a comprehensive Python-based system for macromolecular structure solution. Acta Crystallogr D Biol Crystallogr 66, 213–221, doi:10.1107/S0907444909052925 (2010).

64 Chen, V. B. et al. MolProbity: all-atom structure validation for macromolecular crystallography. Acta Crystallogr D Biol Crystallogr 66, 12–21, doi:10.1107/S0907444909042073 (2010).

