## Supplementary material for "The structural landscape of Microprocessor mediated pri-*let-7* miRNA processing": Supplmentary Information

**Figure S1-**

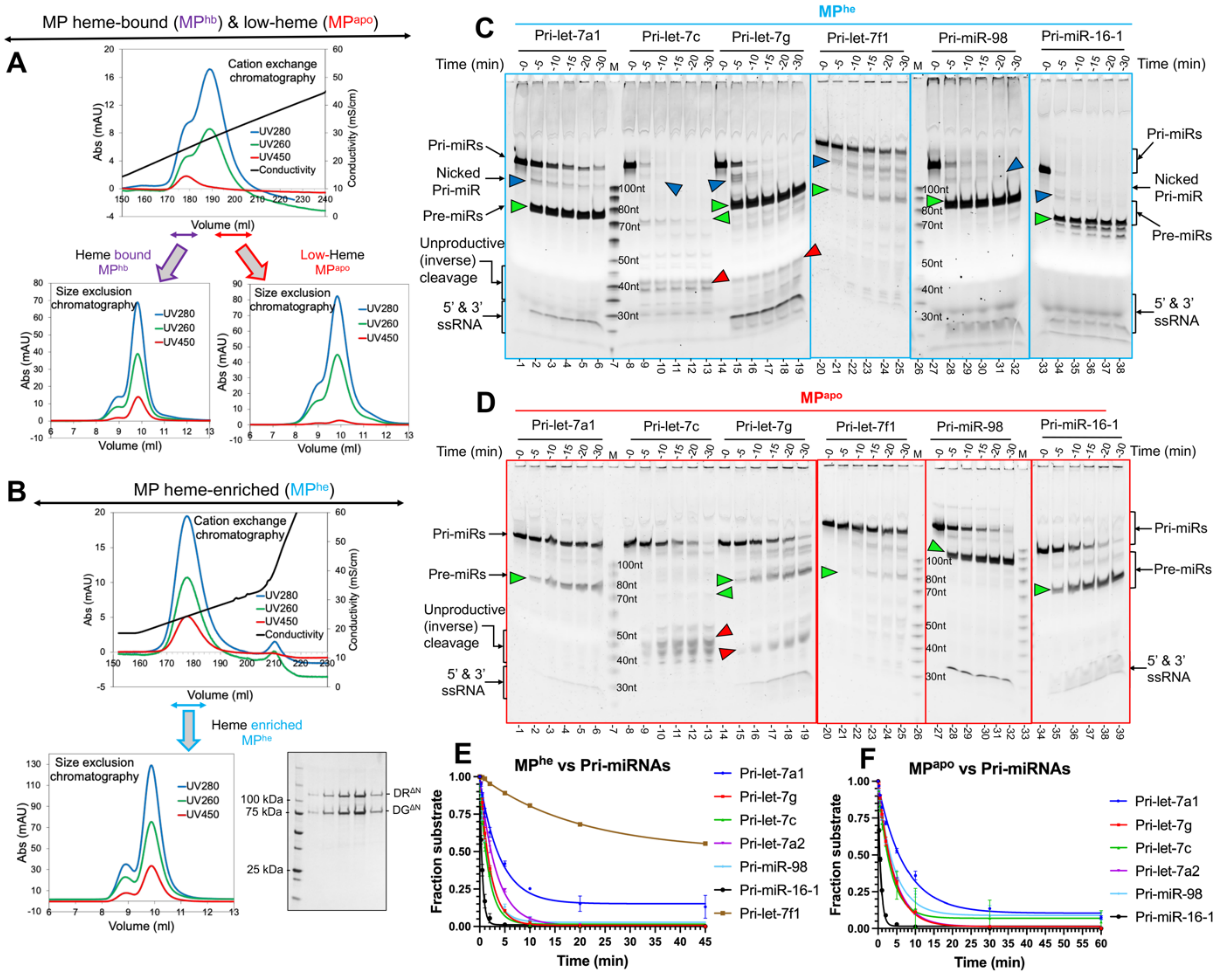

G

| Pri-miRNA | UGUG motif | MP-Heme | $k_{obs}$ (min$^{-1}$) (best fit) | Lower limit (95% CI) | Upper limit (95% CI) | Processing rate relative to Pri-let-7a1 (fold change) |
|---|---|---|---|---|---|---|
| Pri-miR-98 | +ve | enriched | 0.50 | 0.42 | 0.59 | 2.02 |
| Pri-let-7a1 | -ve | enriched | 0.25 | 0.19 | 0.32 | 1.0 |
| Pri-let-7g | -ve | enriched | 0.48 | 0.41 | 0.56 | 1.96 |
| Pri-let-7c | -ve | enriched | 0.57 | 0.46 | 0.70 | 2.31 |
| Pri-let-7a2 | -ve | enriched | 0.31 | 0.28 | 0.34 | 1.25 |
| Pri-let-7f1 | -ve | enriched | 0.05 | 0.04 | 0.05 | 0.20 |
| Pri-miR-16-1 | +ve | enriched | 1.98 | 1.74 | 2.24 | 8.03 |

H

| Pri-miRNA | UGUG motif | MP-Heme | $k_{obs}$ (min$^{-1}$) (best fit) | Lower limit (95% CI) | Upper limit (95% CI) | Processing rate relative to Pri-let-7a1 (fold change) |
|---|---|---|---|---|---|---|
| Pri-miR-98 | +ve | low | 0.23 | 0.16 | 0.34 | 1.64 |
| Pri-let-7a1 | -ve | low | 0.14 | 0.10 | 0.18 | 1.0 |
| Pri-let-7g | -ve | low | 0.25 | 0.22 | 0.28 | 1.78 |
| Pri-let-7c | -ve | low | 0.30 | 0.25 | 0.30 | 2.16 |
| Pri-let-7a2 | -ve | low | 0.23 | 0.20 | 0.27 | 1.67 |
| Pri-miR-16-1 | +ve | low | 1.46 | 1.30 | 1.65 | 10.50 |

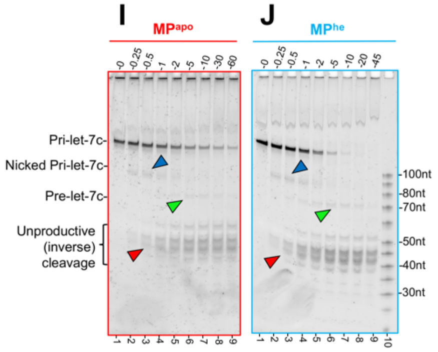

**Figure-S1- MP heme-variants ($MP^{apo}$, $MP^{hb}$ and $MP^{he}$) and *in-vitro* $M^2P^2$ of pri-let-7s.**
(A) Cation exchange chromatogram for MP protein expressed without 5-ALA showing separation of heme-bound MP ($MP^{hb}$) and low-heme MP ($MP^{apo}$) fractions as observed by UV450 absorbance. The separated $MP^{hb}$ and $MP^{apo}$ peaks show homogeneous proteins during size exclusion with average A450/280 ratios of ≤ 0.20 and < 0.10 respectively.
(B) Cation exchange chromatogram of MP expressed with 5-ALA. Heme-enriched MP ($MP^{he}$) eluted as a single peak with an average A450/280 ratio of ≥ 0.25. SDS-PAGE gel showing the purified $MP^{he}$ protein used in this study ($DR^{\Delta N}DG^{\Delta N}$).
Qualitative cleavage assay for different pri-let-7s by (C) $MP^{he}$ and (D) $MP^{apo}$. For most RNAs, $MP^{he}$ generated a clear pre-miRNA (green arrowhead) species in a 30-min time course assay. Pri-let-7c is inversely processed (red arrowhead) by $MP^{he}$ generating a minimal amount of pre-let-7c. $MP^{apo}$ produces significantly less pre-let-7s products compared to $MP^{he}$. Different RNA species including the nicked RNA products (blue arrowhead) are marked. The 100 nt RNA ladder is denoted by “M”.
(E) The plot for the substrate disappearance of different pri-let-7s in near-pre-steady state with $MP^{he}$, (F) $MP^{apo}$. (G) The calculated pri-miRNA cleavage rates with $MP^{he}$. Relative to pri-let-7a1, pri-miR-98 and pri-let-7g are cleaved 2 times faster, while pri-let-7c is inversely processed ~2.3 times faster. The pri-let-7a2 is processed at a similar rate, while pri-let-7f1 and pri-miR-16-1 are cleaved at 0.2 times and ~8 times the pri-let-7a1 rate, by the $MP^{he}$. (H) The calculated pri-miRNA cleavage rates with $MP^{apo}$. The $MP^{apo}$ exhibits slower processing for all pri-miRNAs tested, including pri-miR-16-1 compared to $MP^{he}$, exhibiting heme’s role in $M^2P^2$ of all these non-UGUG motif containing pri-miRs. A representative gel image from the quantitative cleavage assay for pri-let-7c by (I) $MP^{apo}$ and (J) $MP^{he}$ in near pre-steady state conditions. The pre-let-7c (green arrowhead), unproductive cleavage products (red arrowhead) and the nicked product (blue arrowhead) are shown.

**Figure S2-**

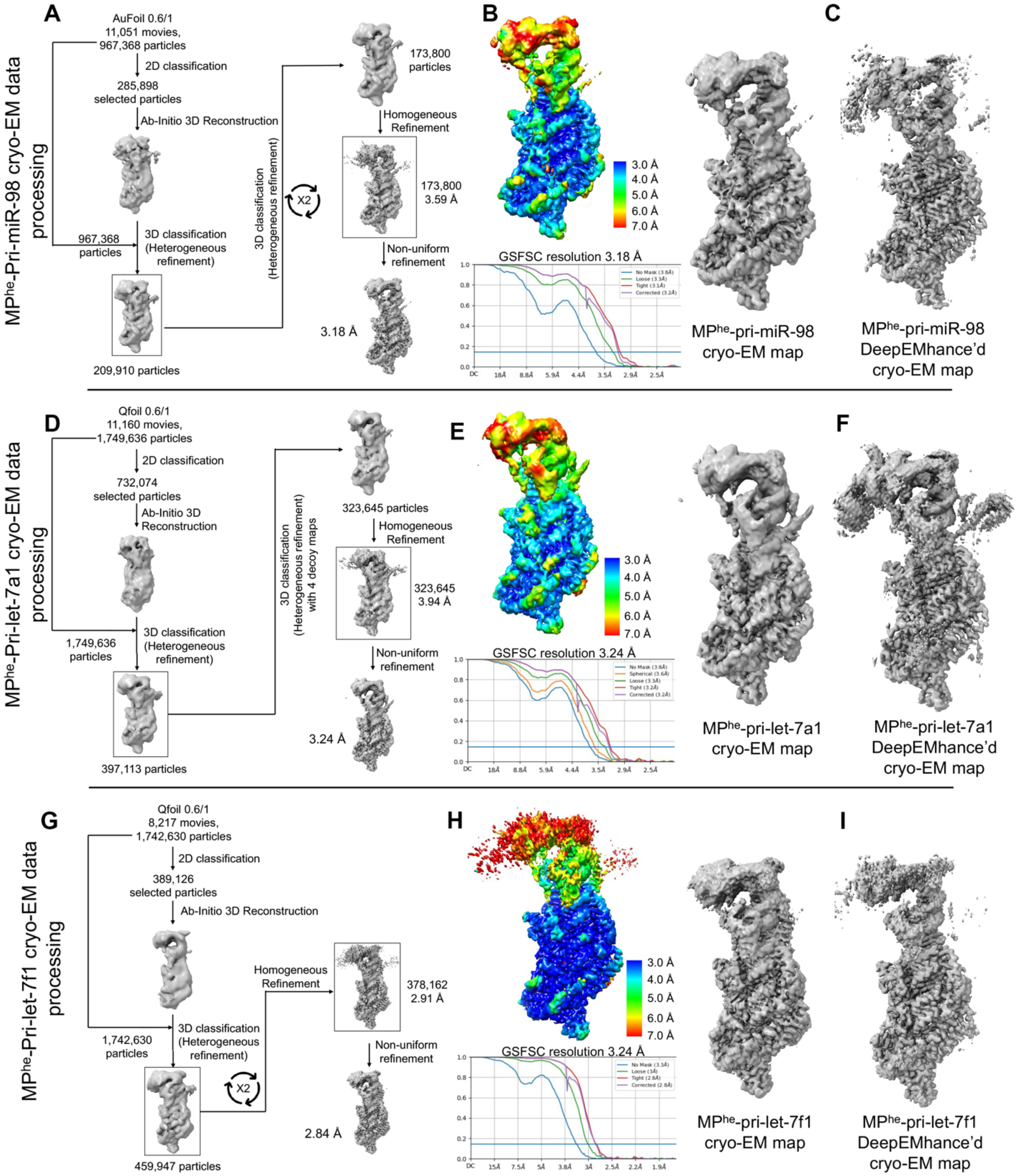
A
MPhe-Pri-miR-98 cryo-EM data processing
AuFoil 0.6/1
11,051 movies,
967,368 particles
2D classification
285,898 selected particles
Ab-Initio 3D Reconstruction
967,368 particles
3D classification (Heterogeneous refinement)
209,910 particles
3D classification (Heterogeneous refinement)
X2
173,800 particles
Homogeneous Refinement
173,800
3.59 Å
Non-uniform refinement
3.18 Å
B
GSFSC resolution 3.18 Å
C
MPhe-pri-miR-98 cryo-EM map
MPhe-pri-miR-98 DeepEMhance'd cryo-EM map
D
MPhe-Pri-let-7a1 cryo-EM data processing
Qfoil 0.6/1
11,160 movies,
1,749,636 particles
2D classification
732,074 selected particles
Ab-Initio 3D Reconstruction
1,749,636 particles
3D classification (Heterogeneous refinement)
397,113 particles
3D classification (Heterogeneous refinement) with 4 decoy maps
323,645 particles
Homogeneous Refinement
323,645
3.94 Å
Non-uniform refinement
3.24 Å
E
GSFSC resolution 3.24 Å
F
MPhe-pri-let-7a1 cryo-EM map
MPhe-pri-let-7a1 DeepEMhance'd cryo-EM map
G
MPhe-Pri-let-7f1 cryo-EM data processing
Qfoil 0.6/1
8,217 movies,
1,742,630 particles
2D classification
389,126 selected particles
Ab-Initio 3D Reconstruction
1,742,630 particles
3D classification (Heterogeneous refinement)
459,947 particles
X2
Homogeneous Refinement
378,162
2.91 Å
Non-uniform refinement
2.84 Å
H
GSFSC resolution 3.24 Å
I
MPhe-pri-let-7f1 cryo-EM map
MPhe-pri-let-7f1 DeepEMhance'd cryo-EM map

**Figure-S2- Cryo-EM structures for class-II pri-let-7s (miR-98, a1 and f1) miRNA in complex with MP$^{he}$.**

The cryo-EM data processing flow chart for (A) MP$^{he}$-pri-miR-98 (D) MP$^{he}$-pri-let-7a1 and (G) MP$^{he}$-pri-let-7f1 structures. Extracted particles were sorted during ab-initio 3D reconstruction and map quality was further improved by iterative rounds of heterogeneous and homogeneous refinement, followed by non-uniform refinement (Punjani et al., 2020). The local resolution map and GSFSC resolution estimate for (B) MP$^{he}$-pri-miR-98 (E) MP$^{he}$-pri-let-7a1 and (H) MP$^{he}$-pri-let-7f1 cryo-EM maps. The Drosha-RNA stem region is resolved to higher resolution than other parts of the complex. DGCR8 dsRBDs are at ~4-5 Å resolution, while HBD is resolved to ~6-7 Å.

Comparisons of the cryo-EM maps processed with and without DeepEMhancer are shown for (C) MP$^{he}$-pri-miR-98 (F) MP$^{he}$-pri-let-7a1 and (I) MP$^{he}$-pri-let-7f1. The overall map density is improved for DeepEMhance'd maps. Both the RNA register and the apical loop position are identifiable in the density-modified maps.

**Figure S3-**

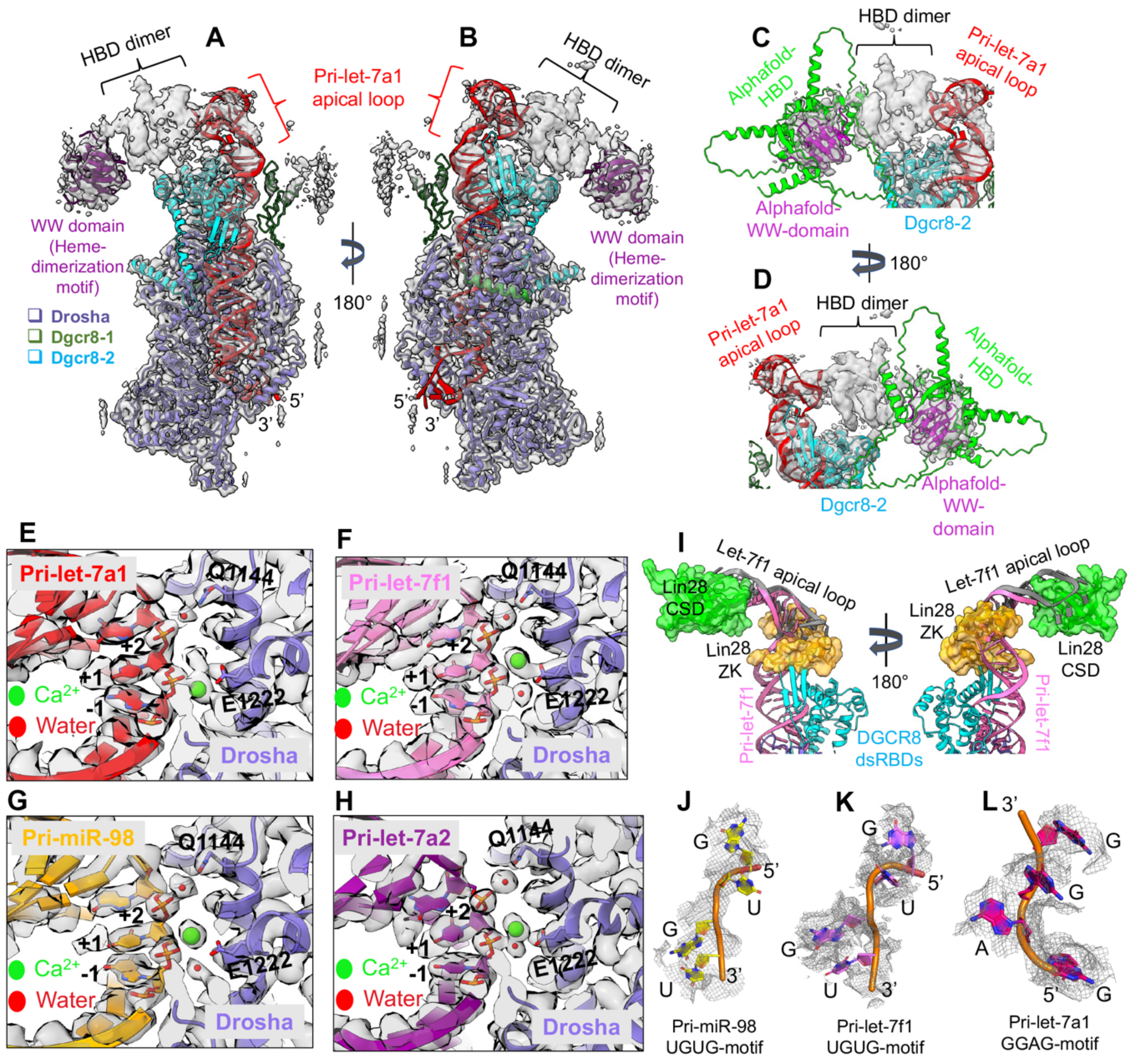
A
B
C
D
HBD dimer
Pri-let-7a1
apical loop
WW domain
(Heme-
dimerization
motif)
Drosha
Dgcr8-1
Dgcr8-2
180°
5'
3'
Alphafold-
HBD
Alphafold-
WW-domain
Dgcr8-2
E
F
G
H
I
J
K
L
Pri-let-7a1
Pri-let-7f1
Pri-miR-98
Pri-let-7a2
Q1144
E1222
+2
+1
-1
Ca2+
Water
Drosha
Let-7f1 apical loop
Lin28
CSD
Lin28
ZK
DGCR8
dsRBDs
Pri-miR-98
UGUG-motif
Pri-let-7f1
UGUG-motif
Pri-let-7a1
GGAG-motif

**Figure-S3- $MP^{he}$-pri-let-7 miRNA complex provides insights into complex assembly and RNA catalysis.**

(A-B) The $MP^{he}$-pri-let-7a1 structure overlaid with the DeepEMhancer modified cryo-EM map, revealing density for the HBD. The WW motif crystal structure (PDBid 3LE4) (purple) is docked into the map. The structural features for the HBD are partially observed but are not modelled in the structure.

(C-D) The AlphaFold-multi (Jumper et al., 2021) predicted HBD model fit into the cryoEM map corresponding to the HBD dimer. Only the WW motif dimer fits reasonably well, but other structural features do not.

The cryo-EM density corresponding to the 5' catalytic site of (E) $MP^{he}$-pri-let-7a1, (F) $MP^{he}$-pri-let-7f1 (G) $MP^{he}$-pri-miR-98 and (H) $MP^{he}$-pri-let-7a2 maps showing the conserved water molecules (red) and $Ca^{2+}$ ion (green). The water coordinating residue Q1144, and $Ca^{2+}$ interacting residue E1222 are shown as sticks. (I) Structural superposition of the $MP^{he}$-pri-let-7f1 apical loop region (pink) with the crystal structure of the Lin28a-let-7f1 apical loop (grey). The zinc-knuckle of Lin28 (shown as a yellow surface) would clash with the DGCR8 dsRBD's (cyan cartoon). The Lin28 CSD is shown as a green surface. (J) The observed cryoEM density for the UGUG motif in pri-miR-98, and (K) pri-let-7f1, and (L) the GGAG-motif in pri-let-7a1.

**Figure S4-**

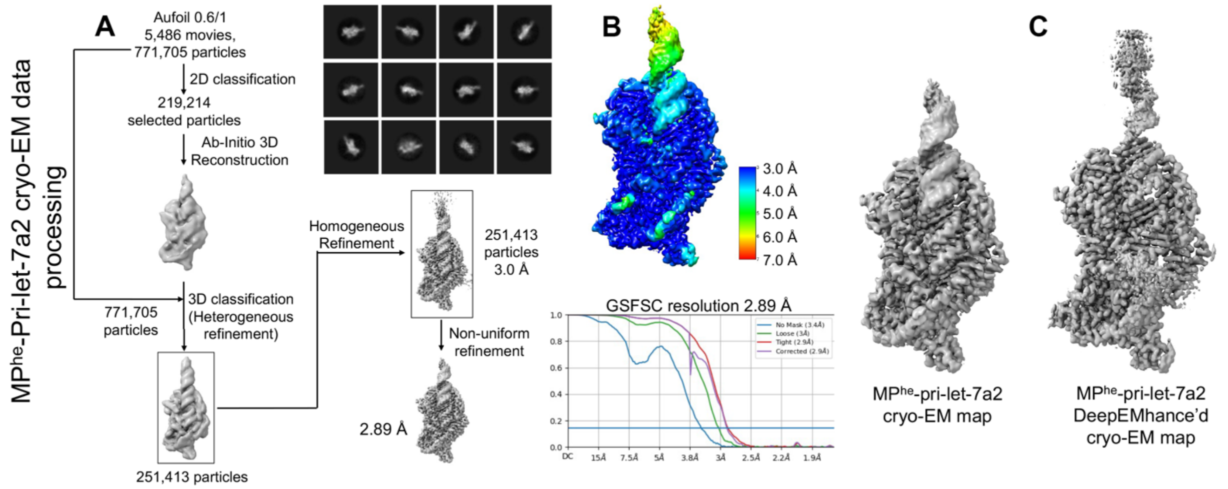

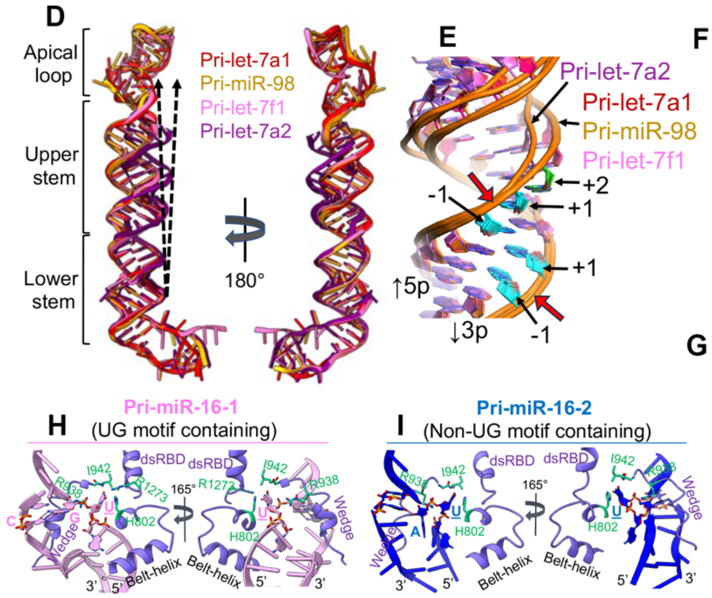

**F**

| Base-pair parameters | | | | | | | | |
|---|---|---|---|---|---|---|---|---|
| Base-pair | Shear (Å) | | | | | | | |
| | Pri-let-7a1 | | Pri-let-7f1 | | Pri-miR-98 | | Pri-let-7a2 | |
| | G-C | -0.20 | G-C | -0.20 | G-U | -1.67 | A-U | 0.04 |
| | A-U | 0.30 | A-U | 0.04 | A-U | 0.11 | G-U | -1.84 |
| **5' catalytic pair** | **G-U** | **-2.1** | **G-U** | **-2.70** | **G-U** | **-2.76** | **U-U** | **-1.10** |
| | A-U | 0.11 | G-C | -0.17 | G-C | -0.15 | U-C | 0.57 |
| 3' catalytic pair | G-C | -0.20 | A-C | -2.20 | G-C | -0.17 | G-C | 0.23 |
| | G-C | -0.16 | G-C | -0.16 | G-C | -0.13 | G-U | -1.13 |

**G**

| Base-pair step parameters | | | | | | | | |
|---|---|---|---|---|---|---|---|---|
| Steps | Rise (Å) | | | | | | | |
| | Pri-let-7a1 | | Pri-let-7f1 | | Pri-miR-98 | | Pri-let-7a2 | |
| | AG/CU | 3.21 | AG/CU | 3.36 | AG/UU | 3.10 | GA/UU | 3.83 |
| | GA/UU | 3.30 | GA/UU | 3.50 | GA/UU | 3.21 | UG/UU | 3.12 |
| **5' catalytic step** | **AG/UU** | **4.82** | **GG/UC** | **4.84** | **GG/UC** | **4.80** | **UU/UC** | **3.30** |
| | GA/UC | 3.45 | AG/CC | 3.40 | GG/CC | 3.52 | GU/CC | 3.40 |
| 3' catalytic step | GG/CC | 3.44 | GA/CC | 3.28 | GG/CC | 3.34 | GG/CU | 3.52 |

**Figure-S4- Structural analysis of class-I and class-II pri-let-7.**

(A) Cryo-EM data processing flow chart for $MP^{he}$-pri-let-7a2 (class-I) complex structure. Selected 2D classes are also shown. (B) Local resolution map and GSFSC resolution estimation plot for $MP^{he}$-pri-let-7a2. (C) Side-by-side comparisons of the cryo-EM map and DeepEMhancer modified map for $MP^{he}$-pri-let-7a2 structure.

(D) Superposition of the class-I and class-II pri-let-7s miRNA bound to $MP^{he}$ in the pre-catalytic state. The positioning of class-I and class-II pri-let-7 upper stems are different, which is indicated by the black arrow. pri-let-7a2 belongs to class-I let7s.

(E) The difference in the backbone geometry in class-I and class-II pri-let-7s, imposed by the 5p +1 bulge nucleotide in 5' cleavage site. The 5p +2 nt (green) is pushed outward in class-II pri-let-7s (7a1, 7f1, miR98) changing the backbone direction. The cleavage sites are marked with red arrows.

The (F) shear and (G) step rise parameters for basepairs around the 5' cleavage site in different pri-let-7s. The basepair involving the 5p +2 nt shows significant changes compared to others (red).

A zoomed-in view of interactions observed for canonical 5' UG motif in pri-miR-16-1 (H) and non-UG motif in pri-miR-16-2 (I). Different residues interacting with RNA nucleotides are shown as sticks (green).

### Figure S5-

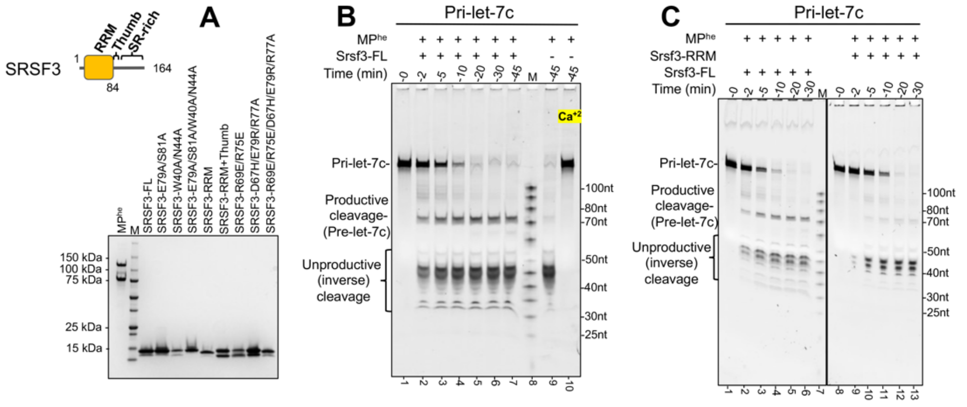

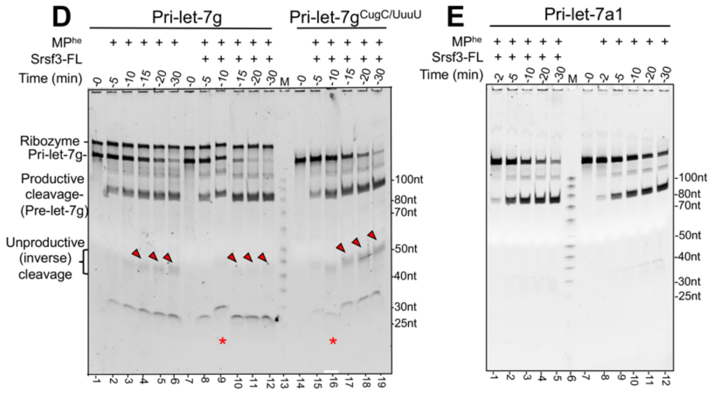

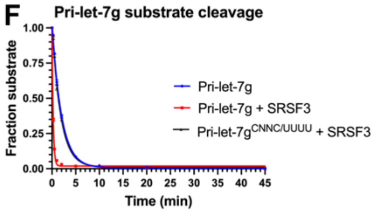

**G**

| | Pri-let-7g | | Pri-let-7g$^{CNNC/UUUU}$ |
|---|---|---|---|
| | No SRSF3 | SRSF3 (650 nM) | SRSF3 (650 nM) |
| $k_{obs}$ ($min^{-1}$) (best fit) | 0.50 | 4.27 | 0.55 |
| Lower limit (95% CI) | 0.44 | 3.85 | 0.48 |
| Upper limit (95% CI) | 0.58 | 4.47 | 0.62 |

**H**

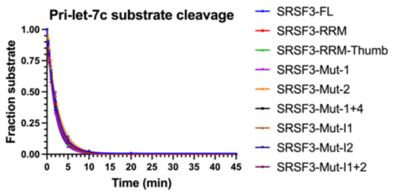

**I**

| Pri-let-7c | Substrate cleavage | | | Inverse processing | | |
|---|---|---|---|---|---|---|
| | $k_{obs}$ ($min^{-1}$) (best fit) | Lower limit (95% CI) | Upper limit (95% CI) | $k_{obs}$ ($min^{-1}$) (best fit) | Lower limit (95% CI) | Upper limit (95% CI) |
| No-SRSF3 | 0.52 | 0.41 | 0.65 | 0.45 | 0.37 | 0.53 |
| SRSF3-FL | 0.44 | 0.42 | 0.45 | 0.41 | 0.35 | 0.47 |
| SRSF3-RRM | 0.46 | 0.44 | 0.47 | 0.41 | 0.37 | 0.45 |
| SRSF3-RRM-thumb | 0.47 | 0.44 | 0.50 | 0.41 | 0.35 | 0.47 |
| SRSF3-Mut-1 | 0.51 | 0.49 | 0.54 | 0.46 | 0.38 | 0.55 |
| SRSF3-Mut-2 | 0.40 | 0.37 | 0.42 | 0.35 | 0.31 | 0.40 |
| SRSF3-Mut-1+2 | 0.48 | 0.46 | 0.50 | 0.48 | 0.35 | 0.65 |
| SRSF3-Mut-I1 | 0.45 | 0.42 | 0.48 | 0.40 | 0.37 | 0.44 |
| SRSF3-Mut-I2 | 0.54 | 0.52 | 0.57 | 0.48 | 0.43 | 0.54 |
| SRSF3-Mut-I1+2 | 0.52 | 0.46 | 0.57 | 0.44 | 0.40 | 0.48 |

**Figure-S5- SRSF3 assists in $M^2P^2$ of CNNC motif containing pri-miRNA.**

(A) SDS-PAGE gel showing purified SRSF3 proteins, mutants/truncations used in this study. SRSF3-E79A/S81A and SRSF3-W40A/N44A correspond to SRSF3-FL with mutations in residues recognizing $C^1$ and $C^4$ nucleotides in the CNNC motif, respectively. SRSF3 RRM and RRM-thumb represent the $SRSF3^{1\text{-}84}$ and $SRSF3^{1\text{-}90}$ truncations. SRSF3-R69E/R75E and SRSF3-D67H/E79R/R77A are SRSF3-FL with mutations in interface-1 and interface-2 of the Drosha-PAZ like domain, respectively. Purified $MP^{he}$ ($DR^{\Delta N}DG^{\Delta N}$) protein is shown in lane 1. The protein ladder is marked as "M".

(B) Processing assay for pri-let-7c in the presence of SRSF3 showing productive cleavage products (pre-let-7c) (also in C). Almost no pre-let-7c is produced in the absence of SRSF3 (lane 9), Addition of 5 mM $Ca^{2+}$ inhibits catalysis. (C) The RRM domain fails to generate pre-let-7c.

(D) Qualitative processing assays of CNNC motif containing pri-let-7g. Urea-PAGE gel analysis shows less unproductive cleavage products (red arrowhead) upon SRSF3 addition (lane-8-12). Pri-let-$7g^{CNNC/UUUU}$ mutant shows a higher level of unproductive cleavage (lanes 15-19). An effect on productive cleavage (pre-let-7g) is not very apparent. The 100nt RNA ladder is marked as "M". Contaminant ribozyme in the pri-let-7g RNA prep is marked and is not catalyzed by $MP^{he}$.

(E) The *in-vitro* RNA cleavage assays showing that catalysis of CNNC-deficient pri-let-7a1 is enhanced in the presence of SRSF3-FL.

(F) The plot for the substrate disappearance for pri-let-7g in near-pre-steady state, showing the effect of SRSF3, and the (G) calculated cleavage rates for each condition.

(H) The plot for the pri-let-7c substrate disappearance in near-pre-steady state, showing the effect of different SRSF3 mutations/truncations, and (I) the calculated substrate cleavage and inverse processing rates in respective condition.

**Figure S6-**

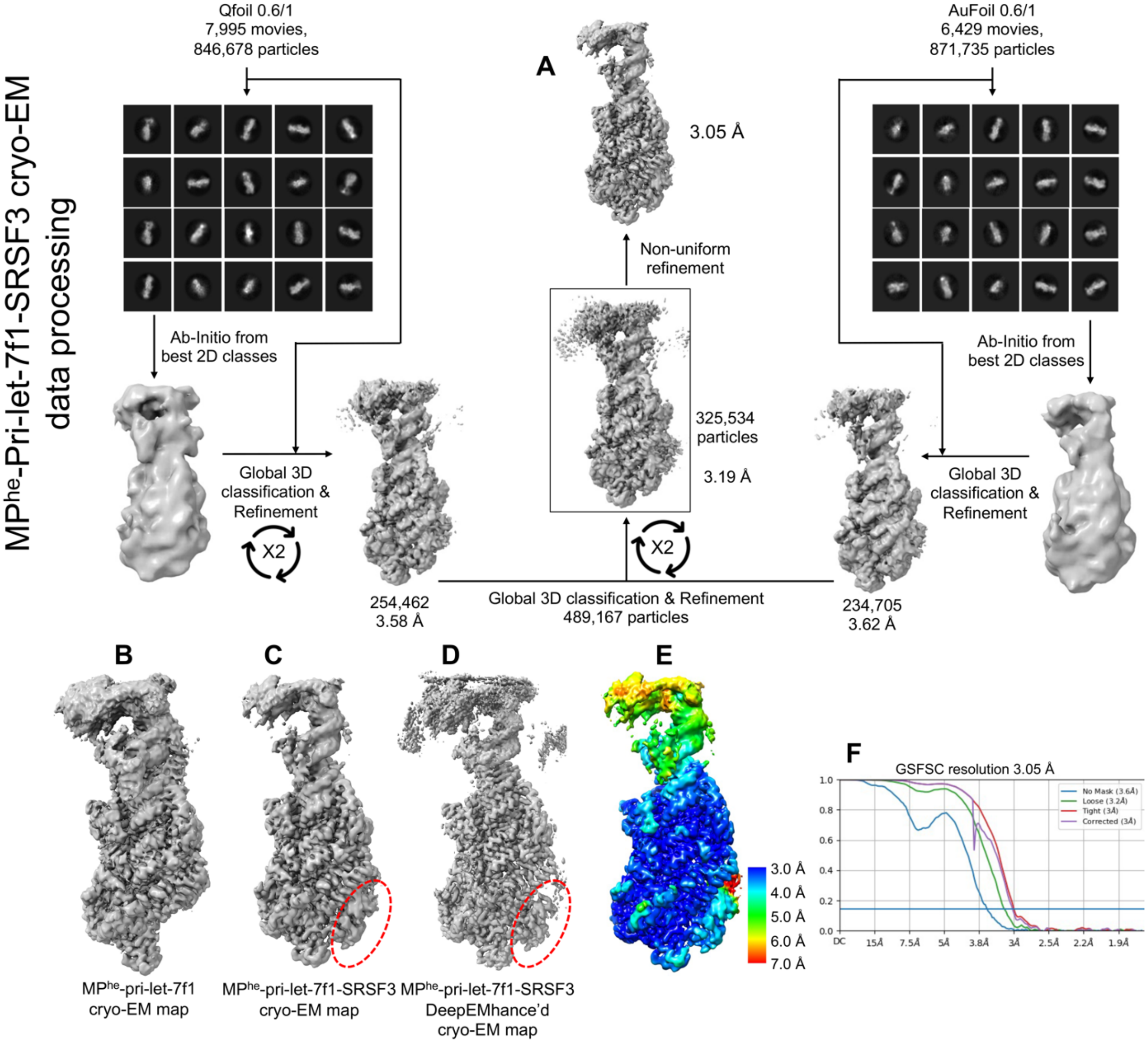
MPhe-Pri-let-7f1-SRSF3 cryo-EM data processing
A
Qfoil 0.6/1
7,995 movies,
846,678 particles
AuFoil 0.6/1
6,429 movies,
871,735 particles
3.05 Å
Non-uniform refinement
Ab-Initio from best 2D classes
Global 3D classification & Refinement
X2
325,534 particles
3.19 Å
254,462
3.58 Å
Global 3D classification & Refinement
489,167 particles
234,705
3.62 Å
B
C
D
E
MPhe-pri-let-7f1 cryo-EM map
MPhe-pri-let-7f1-SRSF3 cryo-EM map
MPhe-pri-let-7f1-SRSF3 DeepEMhance'd cryo-EM map
3.0 Å
4.0 Å
5.0 Å
6.0 Å
7.0 Å
F
GSFSC resolution 3.05 Å
No Mask (3.6Å)
Loose (3.2Å)
Tight (3Å)
Corrected (3Å)
1.0
0.8
0.6
0.4
0.2
0.0
DC
15Å
7.5Å
5Å
3.8Å
3Å
2.5Å
2.2Å
1.9Å

**Figure-S6- The cryoEM structure of MP$^{he}$-pri-let-7f1-SRSF3**

(A) The cryo-EM data processing flow chart for MP$^{he}$-pri-let-7f1-SRSF3 quaternary structure. Two collected cryo-EM datasets were separately processed and 3D heterogeneously filtered particles from both datasets were merged for further processing. The iterative heterogeneous refinement followed by non-uniform refinement resulted in the final map. Selected 2D classes from each dataset are shown.

(B) Cryo-EM map of MP$^{he}$-pri-let-7f1 and (C) MP$^{he}$-pri-let-7f1-SRSF3 show extra density near the Drosha-belt-helix (red dotted oval). The map quality significantly improved after (D) DeepEMhancer processing. (E) Local resolution map and (F) GSFSC resolution estimation plot for MP$^{he}$-pri-let-7f1-SRSF3 map.

**Figure S7-**

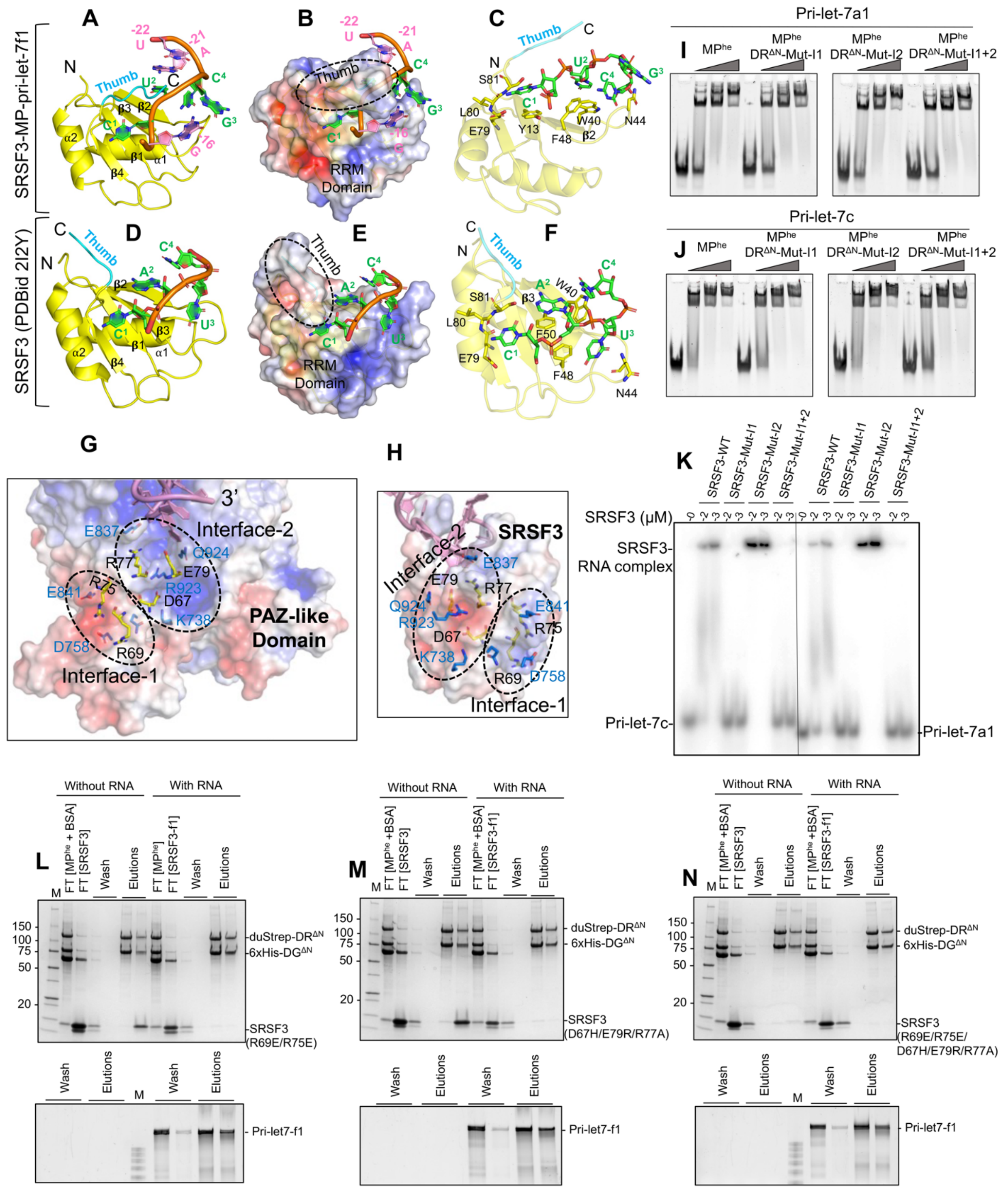
A
B
C
D
E
F
G
H
I
J
K
L
M
N
SRSF3-MP-pri-let-7f1
SRSF3 (PDBid 2I2Y)
Thumb
RRM Domain
Interface-2
Interface-1
PAZ-like Domain
SRSF3
Pri-let-7a1
Pri-let-7c
MPhe
DRΔN-Mut-I1
DRΔN-Mut-I2
DRΔN-Mut-I1+2
SRSF3-WT
SRSF3-Mut-I1
SRSF3-Mut-I2
SRSF3-Mut-I1+2
SRSF3 (μM)
SRSF3-RNA complex
Without RNA
With RNA
Wash
Elutions
duStrep-DRΔN
6xHis-DGΔN
SRSF3 (R69E/R75E)
SRSF3 (D67H/E79R/R77A)
SRSF3 (R69E/R75E/D67H/E79R/R77A)
Pri-let7-f1

**Figure-S7- SRSF3 presents complimentary surfaces for CNNC binding and Drosha PAZ-like domain docking**.

Comparison of the cryo-EM SRSF3-pri-let-7f1 structure with the SRSF3-rCNNC tetranucleotide NMR structure (PDBid 2I2Y). Shown are the cartoon (A & D), electrostatic surface (B & E), and stick (C & F) representations for the respective structures. For the quaternary structure, only 7 terminal nucleotides of the 3p RNA strand are shown. The CNNC motif is colored green. The “thumb” (cyan) is in an open conformation (D-E) and closes on top of the CNNC in the MP-pri-miRNA complex (A-B). The $C^1$, $N^2$, $N^3$ and $C^4$ nucleotides have different interaction patterns in the two structures. The N3 is flipped out in both, and C1 base interactions are conserved. The $C^1$ phosphate is stabilized by Tyr13 in the pri-miRNAs, but is absent in the NMR structure. The $N^2$ base is stacked onto Phe50 from $\beta$3 in NMR structure, but uses Trp40 from $\beta$2 in pri-miRNA. The flipped $N^3$(U) in NMR structure is tucked into a shallow positive groove on the SRSF3 surface, H-bonds with Asn44, with $C^4$ showing no interactions. The flipped $N^3$(G) in the CNNC motif of pri-let-7f1 is sensed by the Drosha belt, with $C^4$ docking into a positive pocket on the SRSF3 surface.

(G) Electrostatic surface of the Drosha PAZ-like domain with interacting SRSF3 residues shown as yellow sticks and interacting residues from Drosha as blue sticks. (H) The electrostatic surface of SRSF3 with interacting residues from the Drosha PAZ-like domain is shown in blue sticks and SRSF3 residues in yellow. Both proteins show complementary positive and negative patches on their surface, which dock onto each other to create the Drosha-SRSF3 interface (black dotted ovals).

Gel-shift assay analyzing the (I) pri-let-7a1 and (J) pri-let-7c binding of $MP^{he}$ with Drosha mutations in SRSF3 interacting interfaces. Interface1 or 2 mutations are marked as $DR^{\Delta N}$-Mut-I1 and $DR^{\Delta N}$-Mut-I2, respectively.

(K) EMSA comparing pri-let-7 binding of SRSF3 with mutations in its Drosha-binding interface. SRSF3 concentrations are marked atop each lane.

(L-N) Pull-down assays of SRSF3 mutants with the $MP^{he}$ and pri-let-7f1. SRSF3-R69E/R75E and SRSF3-D67H/E79R/R77A correspond to residues involved in interactions with Drosha at interface1 and interface2, respectively. SDS-PAGE gel (upper panel) shows that SRSF3 mutants do not co-elute with the $MP^{he}$-pri-let-7f1 complex. Urea-PAGE gel (lower panel) shows the co-eluted pri-let-7f1 stained with SybrGold

#### Supplement Tables:

**Table-S1-** The data collection and model statistics for different MP$^{he}$-pri-let-7 cryo-EM structures.

**Table-S1**: data collection and refinement statistics for cryo-EM structure of MP$^{he}$-pri-let-7s.

| Cryo-EM Map | MP$^{he}$-pri-let-7a1 | MP$^{he}$-pri-miR-98 | MP$^{he}$-pri-let-7f1 | MP$^{he}$-pri-let-7a2 | MP$^{he}$-pri-let-7f1-SRSF3 |
|---|---|---|---|---|---|
| **Data collection and processing** | | | | | |
| Microscope | Titan Krios | | | | |
| Voltage (kV) | 300 | | | | |
| Detector | K3 | | | | |
| Magnification | 81,000x | 81,000x | 105,000x | 105,000x | 105,000x |
| Pixel size (Å) | 1.1 | 1.1 | 0.856 | 0.856 | 0.856 |
| Data collection software | EPU | EPU | EPU | EPU | EPU |
| Defocus range (µm) | 0.6 – 2.2 | 0.7 – 2.2 | 0.6 – 2.2 | 0.6 – 2.0 | 0.7 – 2.2 |
| Total exposure ($e^-/Å^2$) | 72.16 | 42.6 | 77.6 | 78 | 76.6 |
| Frames/exposure | 30 | 30 | 30 | 30 | 30 |
| Exposure/frame ($e^-/Å^2$) | 2.405 | 1.42 | 2.59 | 2.6 | 2.55 |
| Micrographs collected | 11,160 | 11,051 | 8,217 | 5,486 | 7,995 (Q-foil)<br>6,429 (Au-foil) |
| Total extracted particles | 1,749,636 | 967,368 | 1,742,639 | 771,705 | 846,678 (Q-foil)<br>871,735 (Au-foil) |
| Particles for 3D reconstruction | 397,113 | 209,910 | 459,947 | 251,413 | 489,167 |
| Final particles | 323,645 | 173,800 | 378,162 | 251,413 | 325,534 |
| Symmetry | C1 | C1 | C1 | C1 | C1 |
| **Map resolution (unmasked/masked)** | | | | | |
| GSFSC 0.143 | 3.8/3.2 | 3.8/3.2 | 3.3/2.8 | 3.4/2.9 | 3.6/3.0 |
| **Refinement and model validation** | | | | | |
| **Model resolution (unmasked/masked)** | | | | | |
| FSC 0.143 | 3.3/3.2 | 3.2/3.2 | 3.1/3.0 | 3.1/3.0 | 3.2/3.2 |
| FSC 0.5 | 4.5/4.4 | 4.5/4.4 | 4.3/4.2 | 4.2/4.2 | 4.6/4.6 |
| Map CC mask/volume | 0.68/0.71 | 0.70/0.72 | 0.63/0.65 | 0.65/0.67 | 0.60/0.63 |
| Protein residue | 1256 | 1184 | 1180 | 966 | 1257 |
| Nucleic acid | 96 | 97 | 95 | 72 | 100 |
| Water | 1 | 2 | 2 | 2 | 2 |
| **B-factors ($Å^2$)** | | | | | |
| Protein | 180.90 | 178.97 | 178.25 | 154.41 | 167.82 |
| Nucleotide | 76.85 | 81.97 | 80.11 | 83.79 | 78.78 |
| **RMS Deviations** | | | | | |
| Bond length (Å) | 0.004 | 0.005 | 0.003 | 0.002 | 0.003 |
| Bond angle (°) | 0.582 | 0.577 | 0.489 | 0.490 | 0.465 |
| **Ramachandran statistics** | | | | | |
| Favored (%) | 94.86 | 90.63 | 97.17 | 98.23 | 97.34 |
| Allowed (%) | 5.16 | 9.37 | 2.83 | 2.77 | 2.66 |
| Outlier (%) | 0.0 | 0.0 | 0.0 | 0.0 | 0.0 |
| Rotamer outliers (%) | 0.09 | 0.0 | 1.04 | 0.81 | 0.09 |

| | | | | | |
|---|---|---|---|---|---|
| CaBLAM outliers (%) | 2.1 | 1.5 | 2.3 | 1.5 | 2.1 |
| Clash score (all atoms) | 10.51 | 6.14 | 3.96 | 3.65 | 4.84 |
| MolProbity score | 1.90 | 1.57 | 1.34 | 1.15 | 1.37 |
| FSC is Fourier shell correlation, and RMSD is root-mean-square deviation. | | | | | |

**Table-S2-** The sequences for different RNA substrates used in the study. The mutated nucleotides are underlined.

**Table S2. Pri-miRNA sequences used in this study, related to STAR Methods.**

Mutations of wild type (WT) sequences are underlined.

| **RNA name** | **Sequence** |
|---|---|
| Pri-let-7a1: | UGGAUGUUCUCUUCACUGUGGGAUGAGGUAGUAGGUUGUAUAGUUUUAGGGUCACACCCACCACUGGGAGAUAACUAUACAAUCUACUGUCUUUCCUAACGUGAUAGAAAAGUCUGCAU |
| Pri-let-7a2: | UCCAGCCAUUGUGACUGCAUGCUCCCAGGUUGAGGUAGUAGGUUGUAUAGUUUAGAAUUACAUCAAGGGAGAUAACUGUACAGCCUCCUAGCUUUCCUUGGGUCUUGCACUAAACAACAUGGUGAGA |
| Pri-let-7c: | ACAUUGGAAGCUGUGUGCAUCCGGGUUGAGGUAGUAGGUUGUAUGGUUUAGAGUUACACCCUGGGAGUUAACUGUACAACCUUCUAGCUUUCCUUGGAGCACACUUGAGCCGUCGAGGAAUUCUU |
| Pri-let-7f1: | AUUCCAGAAGAAAACAUUGCUCUAUCAGAGUGAGGUAGUAGAUUGUAUAGUUGUGGGGUAGUGAUUUUACCCUGUUCAGGAGAUAACUAUACAAUCUAUUGCCUUCCCUGAGGAGUAGACUUGCUGCAUUAUUUUCU |
| Pri-let-7g: | UUCCUUUUGCCUGAUUCCAGGCUGAGGUAGUAGUUUGUACAGUUUGAGGGUCUAUGAUACCACCCGGUACAGGAGAUAACUGUACAGGCCACUGCCUUGCCAGGAACAGCGCGCCAGCUGCCAAG |
| Pri-miR-98: | AGGAUUCUGCUCAUGCCAGGGUGAGGUAGUAAGUUGUAUUGUUGUGGGGUAGGGAUAUUAGGCCCCAAUUAGAAGAUAACUAUACAACUUACUACUUUCCCUGGUGUGUGGCAUAUUCAGCAU |
| Pri-let-7c$^{CNNC/UUUU}$: | ACAUUGGAAGCUGUGUGCAUCCGGGUUGAGGUAGUAGGUUGUAUGGUUUAGAGUUACACCCUGGGAGUUAACUGUACAACCUUCUAGCUUUCCUUGGAGCACACUUGAGC**UUUU**GAGGAAUUCUU |
| Pri-let-7g$^{CNNC/UUUU}$: | UUCCUUUUGCCUGAUUCCAGGCUGAGGUAGUAGUUUGUACAGUUUGAGGGUCUAUGAUACCACCCGGUACAGGAGAUAACUGUACAGGCCACUGCCUUGCCAGGAACAGCGCGCCAG**UUUU**CAAG |
| Pri-miR-16-1: | AUAGCAAUGUCAGCAGUGCCUUAGCAGCACGUAAAUAUUGGCGUUAAGAUUCUAAAAUUAUCUCCAGUAUUAACUGUGCUGCUGAAGUAAGGUUGACCAUACUCUA |
| Pri-miR-16-2: | CUGACAUACUUGUUCCACUCUAGCAGCACGUAAAUAUUGGCGUAGUGAAAUAUAUAUUAAACACCAAUAUUACUGUGCUGCUUUAGUGUGACAGGGAUACAGCAA |
